# Neural networks representing temporal expectation in mice

**DOI:** 10.1101/2024.04.25.589683

**Authors:** T. Adrian Wendlandt, Patricia Wenk, Julia U. Henschke, Annika Michalek, Toemme Noesselt, Janelle M.P. Pakan, Eike Budinger

## Abstract

The ability to attend to specific moments in time is crucial for survival across species facilitating perception and motor performance by leveraging prior temporal knowledge for predictive processing. Despite its importance, the neural mechanisms underlying the utilization of macro-scale and meso-scale neural resources during temporal processing and their relationship to behavioural strategies and motor responses remain largely unexplored. To investigate the capacity for predictive temporal structure of multisensory stimuli to optimize motor behaviour, we established a behavioural paradigm, in which mice were trained to an auditory-cue and visual-target presented at expected or unexpected temporal delays. Using a combination of stimulus-evoked and resting-state functional magnetic resonance imaging, we examined task-related evoked activity in brain-wide networks and found that that the formation of temporal expectations relying on accumulated sensory information and combined multisensory input involves plasticity across large macro-scale cortical networks comprised of primary sensory systems, sensory association areas including posterior parietal cortex, retrosplenial cortex, prefrontal top-down executive control centres of the brain, as well as hippocampal networks. Additionally, employing in vivo two-photon calcium imaging, we explored local single-cell dynamics within the posterior parietal cortex during this task and found that temporal expectation could be decoded directly from neuronal activity within this brain region. Overall, our study provides insights into the neural correlates underlying the formation of multisensory temporal expectations in the mouse brain and highlights the recruitment of neural resources across temporally-driven statistical learning processes.

## Introduction

Paying attention to particular moments in time is a key cognitive capacity instrumental in the survival of all animals. Since natural environments constantly present us with random and noisy sensory stimulation through which we must interpret behaviourally relevant information, utilising convergent sensory input may enhance perceptual representations (Ernst & Bülthoff, 2004; Henschke et al., 2020; Pakan, Francioni, et al., 2018; Zierul et al., 2017) and optimize adaptive behaviour. The use of temporal information can crucially guide perception and enhance motor performance by utilising prior knowledge about the timing of events for predictive processing (Fiehler et al., 2019; Nobre & van Ede, 2018). As temporal patterns are part of our daily life, these resulting expectations can improve the processing of task-relevant information (Rohenkohl et al., 2012), whereas violations of expectations can also enhance selective attention (M. Tang et al., 2023; X. Tang et al., 2016), and hasten learning (Jin et al., 2020; Jones et al., 2022). It is well established that humans are able to extract regularities in the temporal structure of sensory stimuli (Ball et al., 2022; Ball, Michels, et al., 2018; Nobre & van Ede, 2018), which requires time-keeping mechanisms in the brain (Finnerty et al., 2015). However, how experience encodes temporal expectations across functional neural circuits remains difficult to examine on the circuit-level in humans. Rodent models may prove useful in this regard; however, the fundamental nature of how time is represented in the rodent brain has not been fully established. To date, hippocampus has been identified as a key brain region for temporal processing (Buzsáki & Llinás, 2017), with the discovery of ‘time cells’ that encode successive moments during a delay period (Kraus et al., 2013; Pastalkova et al., 2008) and function in relation to episodic memory (MacDonald et al., 2011; Taxidis et al., 2020).

Beyond the hippocampus, a number of other brain regions have more recently been shown to be important for temporal processing, for example, the entorhinal (Heys & Dombeck, 2018), retrosplenial (Subramanian & Smith, 2024), and motor cortex (Balasubramaniam et al., 2021). In rodents, it has also been shown that knowledge of the timing of auditory stimuli can also improve the speed of motor responses (Jaramillo & Zador, 2011) and that plasticity may be driven by changes in neuronal representation as early as primary sensory cortex (Chapuis & Chadderton, 2018; Huang et al., 2023; Knudstrup et al., 2024; J. Li et al., 2017; R. Li et al., 2023; M. Wang et al., 2020). While temporally predictable unisensory processing may depend on local circuits (Shuler, 2016) and micro-scale cellular processing (Keller & Mrsic-Flogel, 2018; Sharma et al., 2015), the formation of temporal expectations relying on accumulated sensory information and combined multisensory input may require integration via macro-scale cortical networks comprised of wider brain networks (Driver & Noesselt, 2008; Garner & Keller, 2022; Gazzaley & Nobre, 2012; Hawellek et al., 2016; Marek & Dosenbach, 2018; Pinto et al., 2019). Additionally, both human and rodent behaviour is often highly variable, with different behavioural strategies being adopted to accomplish the same goal (Ashwood et al., 2022; Le et al., 2023; Roy et al., 2018). How the use of these macro-scale neural resources during temporal processing may relate to particular behavioural strategies during learning as well as to support the formation of temporal expectations in order to optimize motor responses is unknown.

To investigate this, we established a paradigm to determine the capacity for a predictive temporal structure of multisensory stimuli to optimize motor behaviour and examined task-related evoked activity in brain-wide networks on the macro-scale. Stimulus-evoked (se-fMRI) and resting-state (rs-fMRI) functional MRI are useful techniques to explore these brain-wide circuits and have previously been used to investigate perceptual expectation (Choi et al., 2017; Livesey et al., 2007) and to predict the timing, location, or form of prospective sensory stimuli in humans (Schubotz, 2007). Recently, functional MRI, as a whole-brain approach, has become a powerful method to explore the enhancement of learning performance (Alwashmi et al., 2024) and plasticity in neural networks using graph theory analysis (Ripp et al., 2018; Scharwächter et al., 2022). Here, we apply MRI techniques under multisensory stimulation before and after mice are trained in a paradigm using auditory-cue and visual-target sensory stimuli presented at expected or unexpected temporal delays. We relate behavioural strategies across learning to changes in whole brain networks before and after training, and identify key hippocampal-associated subnetworks that differ in relation to the adopted behavioural strategy. We also found changes in hippocampal-associated and frontoparietal networks that differed according to the capacity for temporal expectation to be readout from motor responses, including activation of the posterior parietal cortex (PPC), an important region for planned movements, attention and sensory association (Lyamzin & Benucci, 2019; Malhotra et al., 2009; Whitlock, 2017). Using in vivo two-photon calcium imaging we investigated local single-cell dynamics within the PPC during this task and found that temporal expectation could also be decoded directly from neuronal activity within this brain region. With this approach, we reveal the neural correlates underlying the formation of multisensory temporal expectations in the mouse brain, and how neural resources are recruited across temporally-driven statistical learning.

## Results

### Behavioural paradigm and formation of temporal expectations in mice

To examine if temporal expectations enhances behavioural motor responses, we established a paradigm in which an expectation about the timing of sensory events is developed through statistical learning, as a target visual stimulus is more likely to be presented at a particular expected (i.e., temporally attended) moment in time following an auditory start cue (Fig. 1A). Beyond their auditory or visual nature, both cue and target stimuli also carry distinct information, with the cue signalling a preparatory signal for motor planning as well as the initiation of ‘time-keeping’, versus the target which acts as a ‘go’ signal for motor execution and the evaluation of predicted results. To implement this task, we first trained mice to lick a reward spout to receive sugar water when a target stimulus was presented (Fig. 1A). This initial training (2-7 days) was conducted with a consistent cue-to-target delay period (2 seconds), where an auditory cue (8 kHz tone) preceded the presentation of a visual target stimulus (checkerboard pattern; Fig. 1A). Following training, mice entered a testing phase (2-5 days, daily sessions) where they were presented with the same auditory-cue and visual-target stimuli but randomly interleaved shorter cue-to-target interval (CTI) periods (0.8 seconds) with longer CTI periods (1.8 seconds). The occurrence of trials with either short or long CTI periods occurred with a fixed probability (75% vs. 25% of trials), which alternated between trial blocks consisting of 100 trials each. Therefore, in each trial block, the only factor that was altered was that a specific temporal delay between the cue and target stimuli occurred more often and, hence, could be predicted with higher likelihood. To probe for temporal expectation we then compared reaction times for two identical delay conditions across blocks (i.e., the average reaction time to the target stimulus during the short CTI when it occurred as the expected condition [75 % likelihood] versus the reaction time to the target stimulus during the short CTI from the unexpected condition [25 % likelihood]; Fig. 1B).

**Figure 1.**
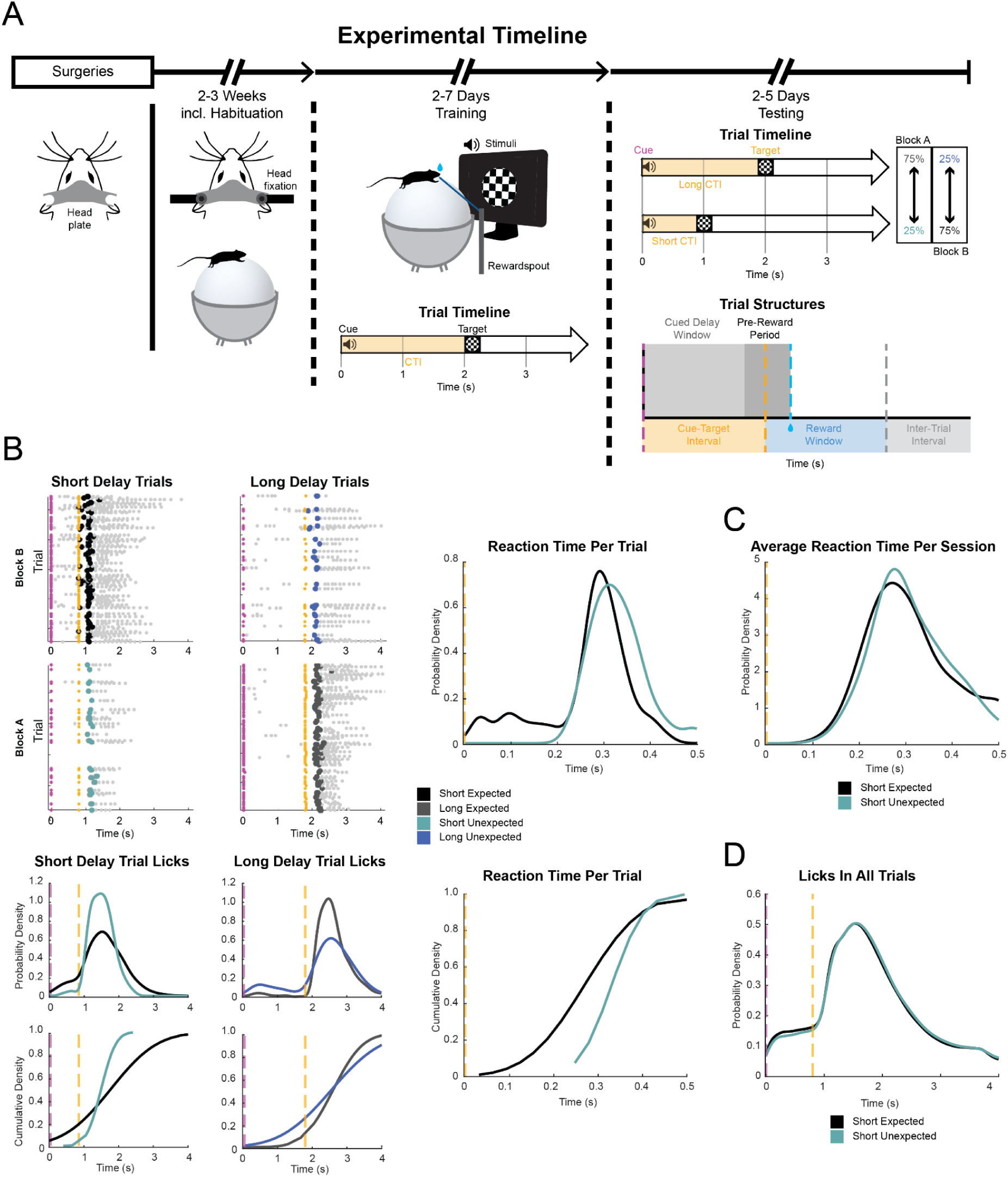
Experimental paradigm and resulting task behaviour indicates the formation of temporal expectations in mice. **(A)** Experimental timeline: 2-3 weeks following headplate attachment mice are trained to lick a reward spout following the presentation of a visual target that occurs after the onset of a 300 ms duration auditory cue (cue-to-target interval [CTI] of 2 s). Following multiple daily training sessions, testing days with either long (1.8 s) or short (0.8 s) CTI delays were presented in blocks, alternating between either 75% or 25% likelihood (Block A or Block B) so that one particular delay period is expected over the course of 100 trials. Black: short expected, blue: long unexpected, teal: short unexpected, grey: long expected. Following the visual target, a 2 s reward window occurred in which licking the reward spout delivered a water reward. This was followed by an intertrial interval (ITI) of 3 s before the start of the next trial. Dashed lines: pink, auditory cue onset; yellow, visual target onset; blue, reward; grey, ITI onset. **(B)** Licking behaviour for one example session. *Top:* raw licking events across blocks. Pink dots: cue onset per trial. Yellow dots: target onset per trial. Grey dots: lick events. The first lick occurring after the target (large coloured dots) was used to calculate reaction time per trial. *Bottom:* probability density and cumulative density (short delay trials: p = 0.0012, Kolmogorov-Smirnov test, n = 719 lick events during expected and n = 132 lick events during unexpected trials; Long delay trials: p = 0.0154, Kolmogorov-Smirnov test, n = 447 lick events during expected and n = 276 lick events during unexpected trials) of all lick events for the example session. *Right*: reaction time probability density and cumulative density (p = 0.046, Kolmogorov-Smirnov test, n = 75 expected and 25 unexpected trials) for the short delay conditions across blocks for the example session. **(C)** Probability density of average reaction times for short expected and short unexpected trials for all sessions (n = 216 sessions). Yellow dashed line, target onset. **(D)** Probability density of all lick events for short expected and short unexpected trials across all sessions (n = 216 sessions). Pink dashed line, cue onset; yellow dashed line, target onset.

A temporal expectation (TE) value (median reaction time for: short unexpected trials - short expected trials) was calculated to quantify the effects of temporal expectation based on the reaction time following the visual target. Long delay trials (1.8 s CTI) are not incorporated into this measure, because a long unexpected target can be predicted due to the omission of the target after the short expected delay period (0.8 s CTI). Hence, a positive TE-value suggests that the animal is attending to the expected moment in time and this has a facilitating effect on motor behaviour, resulting in a faster reaction time (i.e., as measured by the first lick following the presentation of the target stimulus; see probability density distribution of reaction times for an example session Fig. 1B; TE-value: 61 ms; p = 0.0075, Wilcoxon rank sum, n = 75 expected and 25 unexpected trials). Across all sessions, we found a small but significant TE effect with an average reaction time facilitation of 9 ms (p = 0.0074, Wilcoxon sign rank, n= 216, from 49 mice; Fig. 1C). However, we also observed licking behaviour that increased following the presentation of the cue and ramped up in anticipation of the upcoming target (Fig. 1B, D), and this anticipatory licking had a higher probability of occurring in the short delay period (0.8 s following the cue) during the trial blocks when the short delay was expected (see Fig. 1B cumulative distribution, example sessions, p = 0.0012, Kolmogorov-Smirnov test, licks: n = 719 during expected and n = 132 during unexpected trials). Consequently, across the trial period, reaction times could be quicker than the typical reaction time of a mouse (standard rodent reaction time of 200-400ms; (Narayanan & Laubach, 2009; L. Wang & Krauzlis, 2018) and resulted in a bimodal distribution of reaction times across all trials (Suppl. Fig. 1, p = 0.0001, Wilcoxon rank sum, n = 16,091 short expected and 5,417 short unexpected trials, from 216 sessions in 49 mice). These behavioural dynamics may result since, during testing sessions, there were no negative consequences if mice licked at any time outside of the reward window. Therefore, we found that mice were able to utilize the temporal structure of the task to form expectations about the timing of sensory events and to optimize motor behaviour; however, animals could also adopt expectation-independent behavioural strategies to accomplish the task to receive rewards.

### Behavioural strategies and learning-related functional interindividual variability

To further examine the interindividual variability in behavioural strategies between mice over the course of task learning and across sessions of individual mice, the sessions were divided into target-attending and target-invariant behavioural strategies based on if the licking behaviour was spatially confined to the time surrounding the target onset (indicating mice learnt and attended to the significance of the visual target-stimulus) or if licking occurred more constantly throughout the trial (indicating mice did not learn to optimize their behaviour based on the presentation of the visual target stimulus; e.g., Fig. 2A-D, see methods). Note that these two behavioural strategies are not directly related to the capacity for attending to the temporal structure of the task *per se*, as both strategies can result in a positive TE-value based on significantly quicker reaction times during short expected trials (Fig. 2A, Example session 1: 31 ms faster short expected reaction time, p = 0.009, Wilcoxon rank sum, n = 74 expected and 25 unexpected trials; Example session 2: 93 ms faster short expected reaction time, p = 0.005, Wilcoxon rank sum, n = 76 expected and 24 unexpected reaction times). Conversely, both behavioural strategies can also result in a lack of behavioural facilitation of temporal expectation: animals may use a target-attending strategy of attending to the sensory stimuli but not the temporal structure of the task by exclusively waiting for the presentation of the target stimulus before making a response, regardless of the probability of the cue-target delay period; or animals may use a target-invariant strategy but lick with a constant frequency across expected and unexpected trials (see Suppl. Fig. 2).

**Figure 2:**
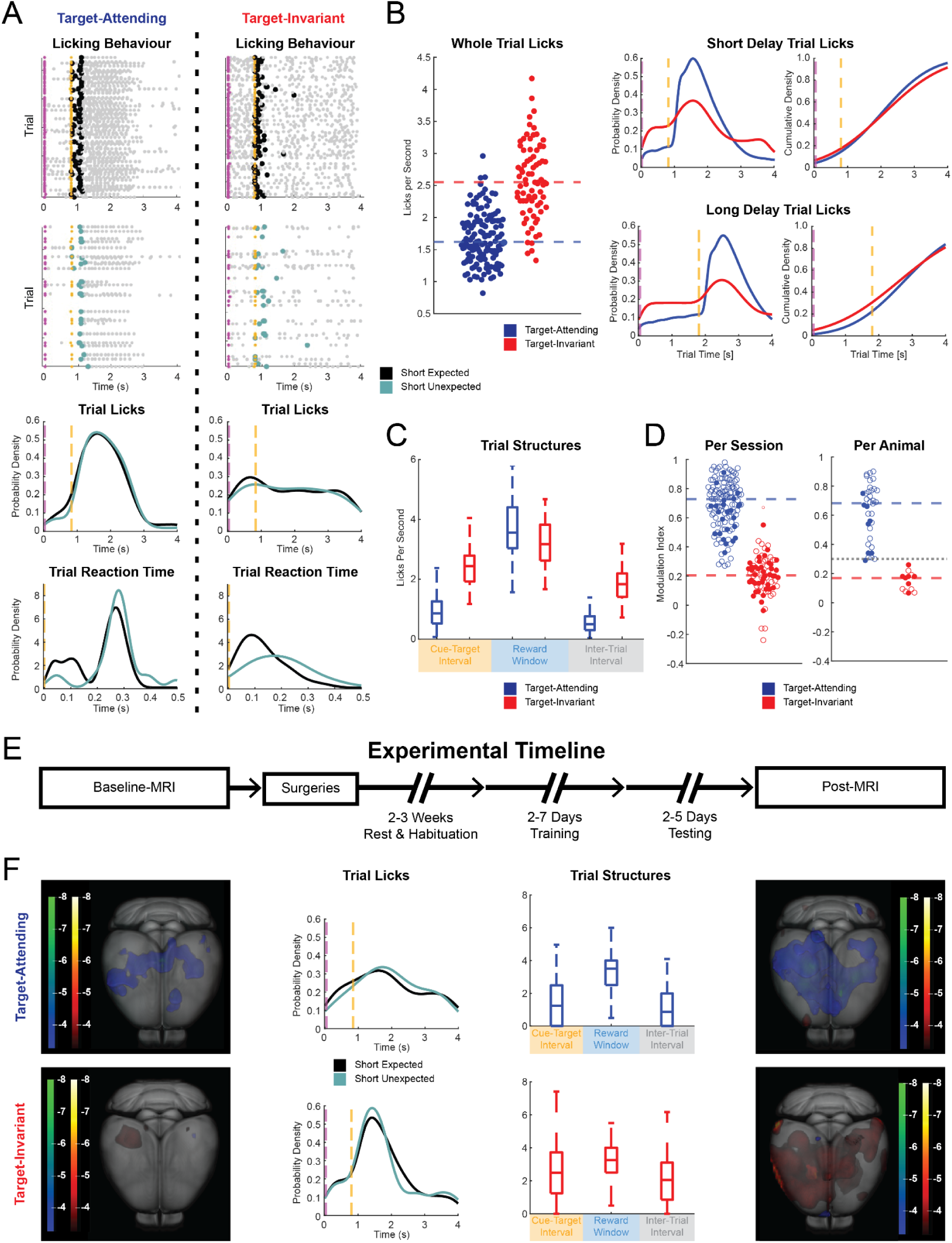
Divergent target-attending and target-invariant behavioural strategies across experimental sessions and animals. **(A)** Representative example sessions of target-attending and target-invariant licking behaviour during short delay trials across blocks. *Top*: raw licking events across blocks. Pink dots: cue onset per trial. Yellow dots: target onset per trial. Grey dots: lick events. The first lick occurring after the target (large coloured dots) was used to calculate reaction time per trial. *Bottom*: probability density of all lick events, and probability density of all reaction times per trial for the corresponding example session. Target-attending example: TE-value: 31 ms, p = 0.009, Wilcoxon rank sum, n = 74 expected and 25 unexpected trials; target-invariant example: TE-value: 93 ms, p = 0.005, Wilcoxon rank sum, n = 76 expected and 24 unexpected trials. **(B)** Evaluation of all target-attentive (red) and target-inattentive (blue) behavioural sessions. *Left*: average lick rate (lick events / s) across all trials shown for each session across groups (p < 0.0001, Wilcoxon rank sum, n = 137 target-attending and 79 target-invariant sessions). *Middle*: probability density of all lick events for sessions in each group during short delay trials and long delay trials. *Right*: cumulative density of all lick events for sessions in each group during short delay trials and long delay trials. Dashed lines: pink, auditory cue onset; yellow, visual target onset. **(C)** Average lick rate (lick events / s) across all trials per session during three trial periods: cue-to-target interval (CTI, yellow), reward window (blue) and inter-trial-interval (ITI, grey) for target-attentive (red) and target-invariant (blue) groups. **(D)** Target modulation index for lick rate averaged across trials for each session (p < 0.0001, Wilcoxon rank sum, n = 137 target-attending and 79 target-invariant sessions) and target modulation index for lick rate averaged across sessions for each animal. Coloured dashed lines indicate median values across groups; grey dashed line: indicates modulation index threshold of 0.3 used to classify mice using overall target-attending and target-invariant behavioural strategies (34 target-attending and 15 target-invariant animals). Filled circles indicate the subset of mice that underwent MRI scans. **(E)** Experimental timeline including baseline MRI scan prior to behavioural training and post training MRI scan following training. **(F)** BOLD activation pattern of one example target-attending and one example target-invariant mouse at baseline and post training fMRI scan. A general linear model approach (p < 0.001, no correction for multiple comparison) was used for statistical analysis of the signal change upon repetitive visual stimulation. The heatmap is based on the resulting t-values (a t-value of 3.29 equals p < 0.001), with brighter colours indicating higher statistical significance. Blue-green colour scale represents negative BOLD signal change and red-yellow colour scale positive BOLD signal change. Probability density is shown for all licks for trials with short expected and short unexpected delays for an individual session of the respective example mice. Pink dashed line, cue onset; yellow dashed line, target onset. Average lick rate (lick events / s) averaged across all trials for different trial periods is also shown for the individual session of the respective mice, during: the cue-to-target interval (CTI, yellow), reward window (blue) and inter-trial-interval (ITI, grey). In both examples, the trial structure significantly predicts licks per second, but more variance in the data can be explained for the target-attending example (target-attending: F_(2, 297)_ = 88.88, p < 0.0001, adjusted R^2^: 0.36; target-invariant: F_(2,297)_ = 9.96, p< 0.0001, adjusted R^2^: 0.05; one-way Anova, constrained (type III), sums of squares).

Since the strategy of constant licking throughout the trial can make defining a specific reaction time following the target onset an unreliable measure of temporal expectation, we also quantified the rate of licking across various periods of the trial structure as a readout (Fig. 2B-D). Target-invariant sessions have a significantly higher number of licks when averaged across the whole trial period (p < 0.0001, Wilcoxon rank sum, n = 137 target-attending sessions and 79 target-invariant sessions; Fig. 2B). Differences in the pattern of licking behaviour across groups are evident when comparing the probability and cumulative density of licks for trials with both long and short CTI delays (Fig. 2B), with significant main effects for behavioural strategy as well as trial structure (Fig. 2C; strategy: *F_(1,642)_* = 198.3, p < 0.0001; structure: *F_(2,642)_* = 616.2, p < 0.0001; two-way mixed-effects ANOVA, constrained (type III), sums of squares). Whereas target-invariant strategies result in a significantly higher lick rate in both the CTI and ITI periods (CTI: p < 0.0001, 95% C.I. = [-1.79, -1.25]; ITI: p < 0.0001, 95% C.I. = [-1.26, -0.99]; Tukey-Kramer correction), target-attending strategies had a higher lick rate specifically following the target onset (reward window: p < 0.0001, 95% C.I. = [0.20, 0.74]; Tukey-Kramer correction). Calculating a lick modulation index (average licks/s in 1 s window: pre-target - post-target / pre-target + post-target; Fig. 2D) allowed us to evaluate the impact of target presentation on the rate of licking behaviour across sessions as well as to classify the behavioural strategy across individual animals. Animals were considered as using an overall target-attending strategy, if the average lick rate following target onset was at least doubled (i.e., average lick modulation index across sessions ≥0.3; Fig. 2D).

### Stimulus-evoked fMRI BOLD responses with behavioural strategy before and after training

To determine the underlying neural resources that support the different behavioural strategies, stimulus-evoked BOLD (blood-oxygen-level-dependent) signal change was investigated both before and after behavioural training (Fig. 2E) in a subset of mice considered to be using target-attending versus target-invariant behavioural strategies according to this classification (Fig. 2D; n = 15 mice, 8 target-attending and 7 target-invariant). Lightly sedated animals were placed in a high-field (9.4 T) small-animal MRI scanner and presented with the similar sensory stimuli as used in the behavioural training, i.e., 8 kHz tone and checkerboard pattern visual stimulation. Overall stimulus-evoked BOLD responses to the visual stimulus of a representative mouse using either the target-attending or target-invariant behavioural strategy are illustrated in Figure 2F, also in relation to the behavioural performance of the animals. Using a general linear model approach to assess whole-brain patterns of BOLD responses during se-fMRI, mice using either behavioural strategy already showed significant differences in the BOLD patterns across the brain at baseline; following training, these differences were even more pronounced. Generally, we found mainly positive BOLD responses for the target-invariant mice, even after training, which indicates a higher demand of neural resources to process the temporal structure of sensory stimuli. In contrast, target-attending mice showed more negative BOLD responses suggesting that they recruit less neural resources (see also discussion).

To further examine differences in whole-brain activation across at the group level, we calculated BOLD difference maps (target-attending - target-invariant mice) for auditory (cue stimulus), visual (target stimulus), and combined audio-visual (cue+target stimulation) se-fMRI at baseline (Fig. 3A) and post training (Fig. 3B). Again, target-attending and target-invariant mice showed clear differences (for regionally individual t-values based on a p-value < 0.001 see Suppl. Tab. 2) in the resulting BOLD contrast to the three stimulus categories already at baseline. Target-invariant mice showed higher BOLD contrast, i.e., a more negative contrast, than target-attending animals in many brain regions, for example, in regions involved in sensory processing (e.g., visual areas [VIS], SC, inferior colliculi [IC], caudoputamen [CP]), executive functions (prelimbic area [PL], cingulate area [CG]) and learning and memory task-related regions (hippocampal formation [HPF], in particular CA1/3 regions, subiculum [SUB] and entorhinal area [ENT]) (Fig. 3A,C,D). On the other hand, the somatosensory cortex (SSCX) and SC showed more positive contrast in target-attentive mice (Fig. 3A,C,D). These findings indicate that there are pre-existing differences in hemodynamic changes across brain regions associated with sensory processing as well as learning and memory for the mice that would later adopt a target-attending versus target-invariant behavioural strategy during the paradigm.

**Figure 3.**
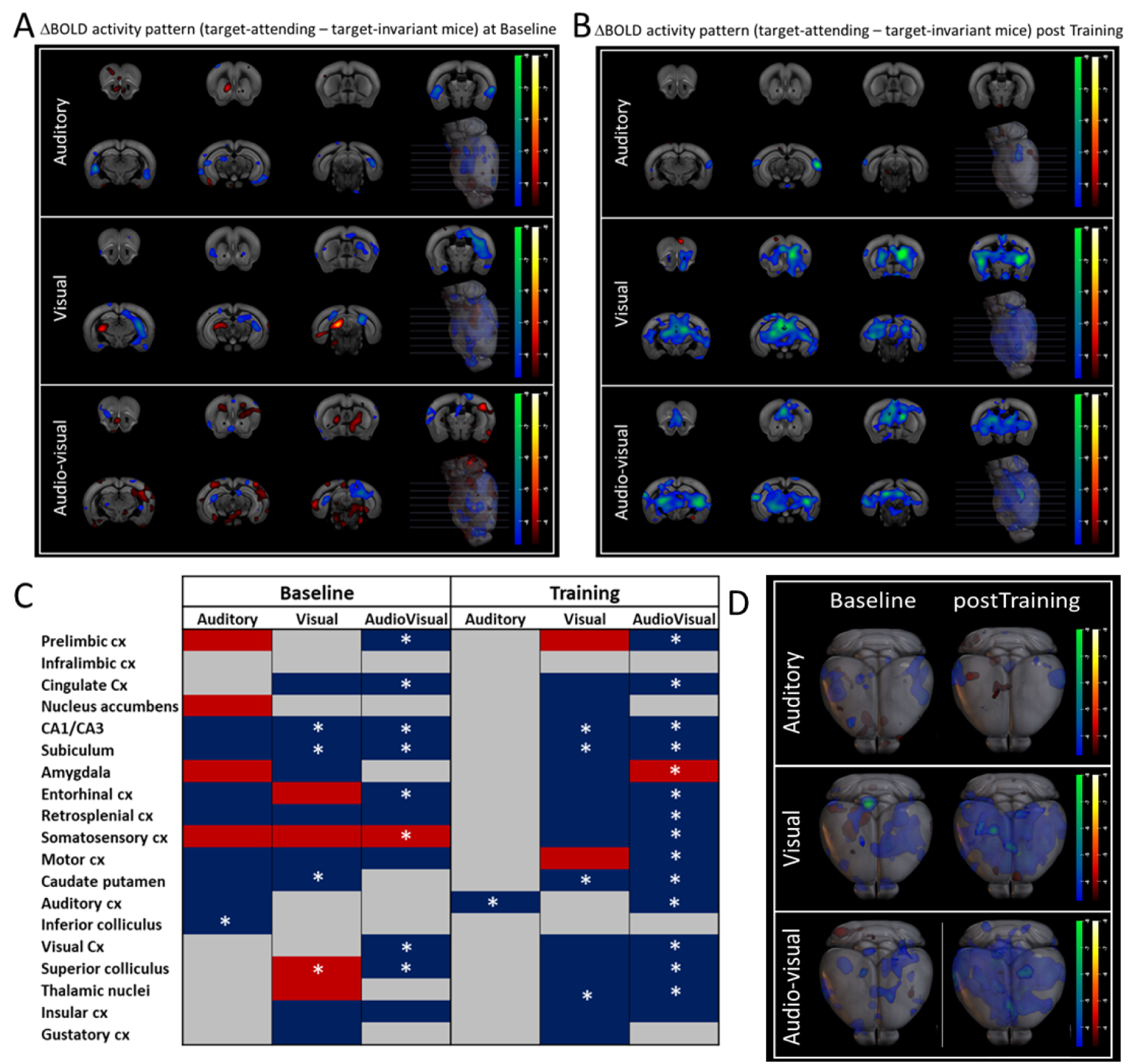
Quantification of whole-brain BOLD activity patterns for target-attending and target-invariant mice at baseline and post training. Difference maps of BOLD activity patterns for all three stimulation paradigms (auditory [cue], visual [target], audio-visual [simultaneous cue+target]) for mice with a target-attending (n=7) behavioural strategy minus mice with a target-invariant (n=7) behavioural strategy at baseline **(A)** and mice with a target-attending (n=8) behavioural strategy minus mice with a target-invariant (n=7) behavioural strategy post training **(B)**. The significance level for the individual maps was set to p < 0.001 (t-value of 3.29) before analysis. The heatmap is based on the resultant t-values before subtraction of the images, here brighter colours indicate higher differences between the two maps. The blue-green colour scale represents negative differences and the red-yellow colour scale positive differences between the BOLD contrast of groups (target-attending - target-inattentive). **(C)** Observed differences of brain areas with detected differences for all three stimulation paradigms individually with their corresponding appearance (blue for negative difference [indicating that target-inattentive mice had higher BOLD response], red for positive difference [indicating that target-attending mice had higher BOLD response], grey marks undetectable regions). An asterisk highlights regions with statistical significant differences based on a two-sample unpaired t-test (p < 0.001). Individual t-values are listed in Supplementary Tables 2 and 3. **(D)** 3D volume rendering highlights the whole-brain differences with the same colour bars as shown in (A).

After training, an even larger number of brain regions showed a negative BOLD contrast, i.e., overall higher activation in target-invariant and lower activation in target-attending mice (Fig. 3B,C,D). This was particularly the case for visual and audio-visual stimulation. In response to the visual target, we found significantly decreased BOLD responses, for regionally individual t-values based on a p-value < 0.001 (see Suppl. Tab. 3, for the target-attending group) in hippocampal (e.g. CA1/CA3, t-value of -6.64 for audio-visual stimulation, and SUB), striatal (CP, t-value = -4.88 for visual stimulation) and thalamic regions, indicated by the decreased BOLD contrast (Fig. 3B). Motor (MoCx) and prelimbic regions (PL) showed increased activity patterns in the target-attending mice. During combined audio-visual stimulation, in addition to regions which were already evident at baseline (i.e., PL/CG, CA1/3/SUB/ENT, VIS/SC) many more regions showed a significant negative BOLD contrast. Among those are primary sensory areas (auditory, visual, somatosensory) as well as the retrosplenial cortex (RSC, t-value = -3.45 for audio-visual stimulation), a brain region that is both highly connected with the hippocampus and involved in context-specific learning and memory (Fig. 3B,C) (Sun et al., 2021; Todd et al., 2019). Interestingly, the amygdala (AMY) showed a significant positive BOLD contrast during audio-visual stimulation which might support their role in triggering temporal prediction errors (Tavares et al., 2022).

Altogether, differences in BOLD activity patterns both before and after training occur across diverse brain networks between mice adopting a target-attending or a target-invariant behavioural strategy during the task, but are particularly prominent in areas involved in processing task-relevant information across learning and differ more significantly after training in the presence of the visually-evoked target stimulus. This is in line with divergence of their behavioural strategy, where attention is differentially directed towards the visual target during learning and may represent the use of fundamentally divergent functional networks.

### Functional connectivity patterns at rest across mice with divergent behavioural strategies

Network dynamics and task-related changes before and after training can further be defined based on metrics of functional connectivity (Biswal et al., 1995; Ma et al., 2016; Scharwächter et al., 2022). These measures relate to the temporal dependence between spatially remote neurophysiological events, with positive correlations reflecting integration or temporally coordinated activity across brain regions. We used functional connectivity calculated from rs-fMRI to identify connectivity patterns of the target-attending and target-invariant groups of mice and observe task-related brain network changes before and after training. We examined the correlation of 49 brain regions bilaterally segregated across hemispheres (98 total regions, see Suppl. Tab. 1; Fig. 4A). While correlations generally decreased in the target-invariant group after training (Fig. 4A,B; for mean correlation values ± s.e.m. see Suppl. Fig. 4), we found clear increased correlations in the target-attending group following training (Fig. 4A,B), indicating a global increase in functional connectivity specific to mice that adopted this behavioural strategy (mean correlation value ± s.e.m. for target-attending mice at baseline/ post training = 0.024 ± 4.9E-5/ 0.055 ± 7.3E-5). A graph theory approach was then applied to calculate regional connectivity in a specific subnetwork (Scharwächter et al., 2022). Here, we focused on a subnetwork of relevant sensory, prefrontal, and memory-related brain areas (nodes) that also showed activation during the se-fMRI (Fig. 4C, see also Fig. 3) and examine their structural and functional relation (edges). This approach recapitulates the differences between these two groups of mice at baseline, in line with our se-fMRI findings (Fig. 3). Mice that later adopted a target-attending behavioural strategy tend to have more connections between nodes with a higher average node degree within this subnetwork (i.e., the sum of all edge weights connected to a node; for the mean node degree ± s.e.m. see Suppl. Fig. 6), particularly in relation to the CG, RSC, VIS and PPC (Fig. 4C). After training, we found changes in the functional connectivity patterns within this subnetwork of both groups, with both groups showing high levels of functional connectivity within the subnetwork following training. The most dominant increase of node degree was visible for the HPF for the target-attending mice with a mean node degree of 14.43 ± 4.29 at baseline and 20.19 ± 4.63 after training (Fig. 4C). In addition, the CG, PPC and RSC remained with a higher node strength for the target-attending group. Conversely, an increased node degree was observed for the target-invariant group in parahippocampal regions and the temporal association area (TEa). Here, the node degree of TEa increased from 8.00 ± 3.47 to 11.29 ± 5.70, whereas for target-attending mice it decreased from 8.36 ± 3.19 to 4.00 ± 1.93 after training. This indicates an aligned node degree at baseline with a discriminative development due to training. Note, the node strength of auditory areas remained unchanged across groups and following training, in line with the focus on visual target dynamics defining the behavioural strategies during learning across these groups.

**Figure 4.**
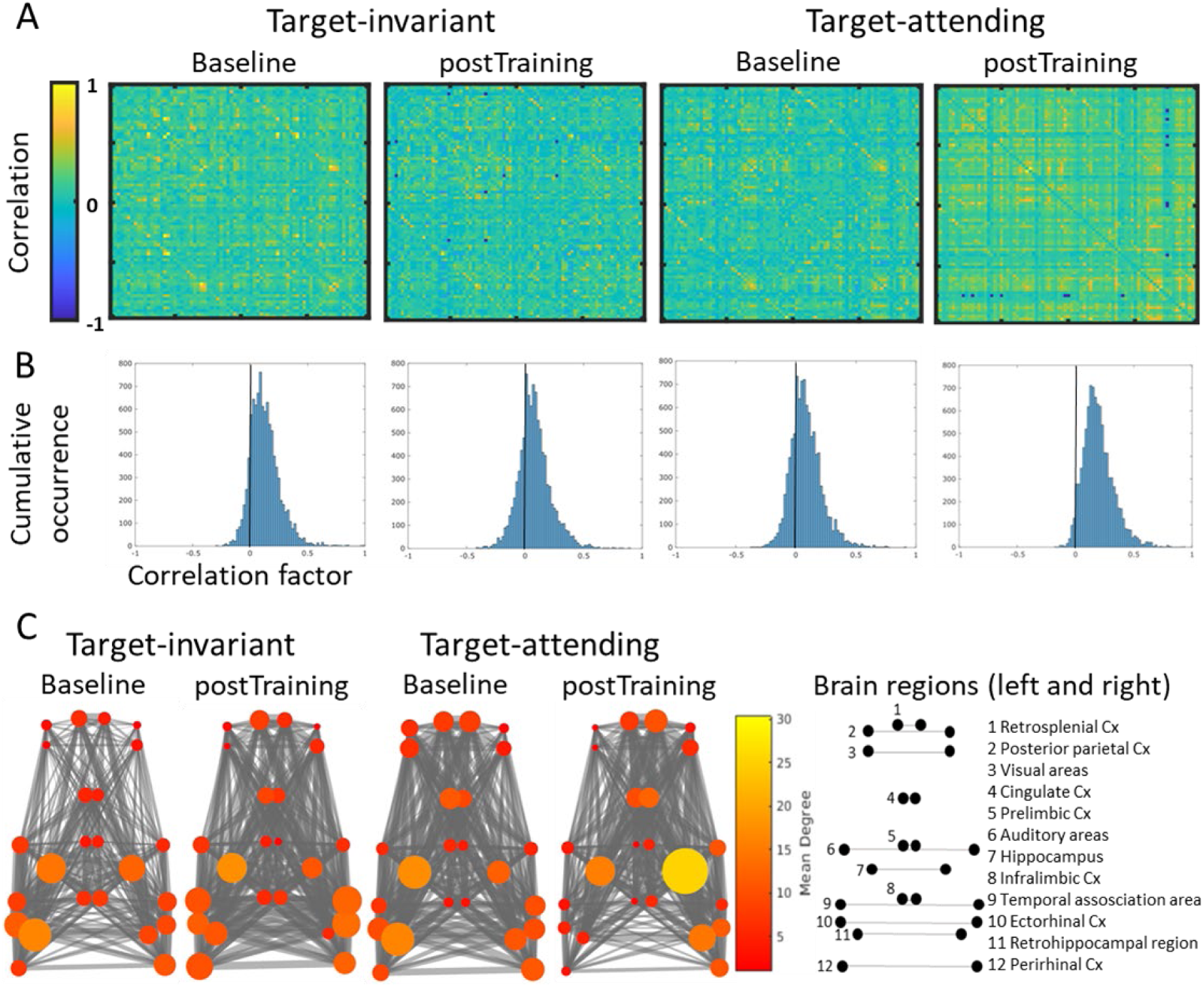
Functional connectivity of resting-state fMRI (rs-fMRI) before and after training for mice with target-attending versus target-invariant behavioural strategies. **(A)** Correlation matrices across 49 brain regions bilaterally (98 total areas; see Suppl. Tab. 1) for target-invariant (n = 8) and target-attending (n = 7) mice calculated for baseline and post training rs-fMRI scans. Heatmap indicates the correlation strength based on the z-transform of the Pearson correlation coefficient (mean correlation value ± s.e.m. for target-invariant at baseline = 0.033 ± 6.4E-5 and post training 0.026 ± 5.1E-5; mean correlation value ± s.e.m. for target-attending at baseline and post training = 0.024 ± 4.9E-5 and 0.055 ± 7.3E-5, see Suppl. Figure 4). **(B)** Histogram of Pearson correlation coefficients across all 98 brain areas as calculated in (A). **(C)** A graph theory approach for selected brain areas of interest is shown across networks of interest (numbered list on right) for target-invariant and target-attending mice at baseline and post training rs-fMRI scans. Size and colour of the nodes represent the node degree (larger and brighter colour represents higher node degree). Mean node degree values ± s.e.m. of presented regions are available at Supplementary Figure 6.

### Efficient use of temporal expectation to optimize behaviour and related functional networks

As previously noted, while target-attending versus target-invariant behavioural strategies can explain dynamics specific to the interindividual variability based on learning of the task parameters, they are not directly related to the capacity for attending to the temporal structure of the task *per se*, as both strategies can result in a positive TE-value (Fig. 2, Suppl. Fig. 2). To further examine the network dynamics underlying temporal processing specifically, we quantified behaviour that is facilitated by attending to the temporal structure of the task and the strategic use of temporal expectations to optimize motor behaviour. While temporal expectation can be measured directly based on the reaction time dynamics following the short delay period and the related TE-value (Fig. 1) (Ball, Michels, et al., 2018; Stefanics et al., 2010), various other task dynamics can also be considered (Fig. 5A): i) if a long delay is expected, occurrence of the target at an earlier time point is *surprising* and should result in a slower reaction time; ii) cue onset facilitates time-keeping for expectation-based behaviour, hence, the frequency of *anticipatory* licking should increase within the cue-target interval period when the short delay is expected; iii) omission of an early expected stimulus primes behaviour for the occurrence of the stimulus at a later time point and should result in a *priming* effect of anticipatory licking directly before the reward is given. Across all sessions, we found evidence of temporal expectation that fits all three of these criteria (Fig. 5A); however, again with substantial interindividual variability. We classified mice along these three dimensions (Fig. 5B) to identify sessions in which mice attended to the temporal structure of the task and showed strong evidence for the formation of temporal expectations (time-attending) versus when they showed no, or only weak, evidence of the formation of temporal expectations to optimize behaviour (time-invariant). Using k-means clustering and the Euclidean distance between sessions to define two clusters along these dimensions, we then evaluated the resulting TE-value of these two groups to determine if the three separate criteria of temporal expectation also reduce interindividual variability as measured by the session TE-value. Indeed, we found that the time-attending cluster had a significantly higher TE-value than the time-invariant cluster (time-attending: 28 ± 5 ms; time-invariant: -8 ± 7 ms; p < 0.0001, Wilcoxon rank sum, n = 112 time-attending sessions and 104 time-invariant sessions; Fig. 5B). To classify how each individual animal approached the task and their overall use of temporal attention to optimize behaviour, we first averaged these metrics across the sessions for each mouse and then performed the same clustering methods and found similar dynamics in the TE-values across these groups on the level of individual animals (time-attending: 13 ± 2 ms; time-invariant: -4 ± 4 ms; p = 0.0001, Wilcoxon rank sum, n = 27 time-attending mice and 22 time-invariant mice; Fig. 5C). These clustered groups were then used to examine the underlying neural resources that support the use of task-related temporal attention by evaluating the stimulus-evoked BOLD response both before and after behavioural training in the same subset of mice as above (see Fig. 3, 4), but distributed into time-attending versus time-invariant groups (Fig 5C; n = 10 time-attending mice and 5 time-invariant mice).

**Figure 5:**
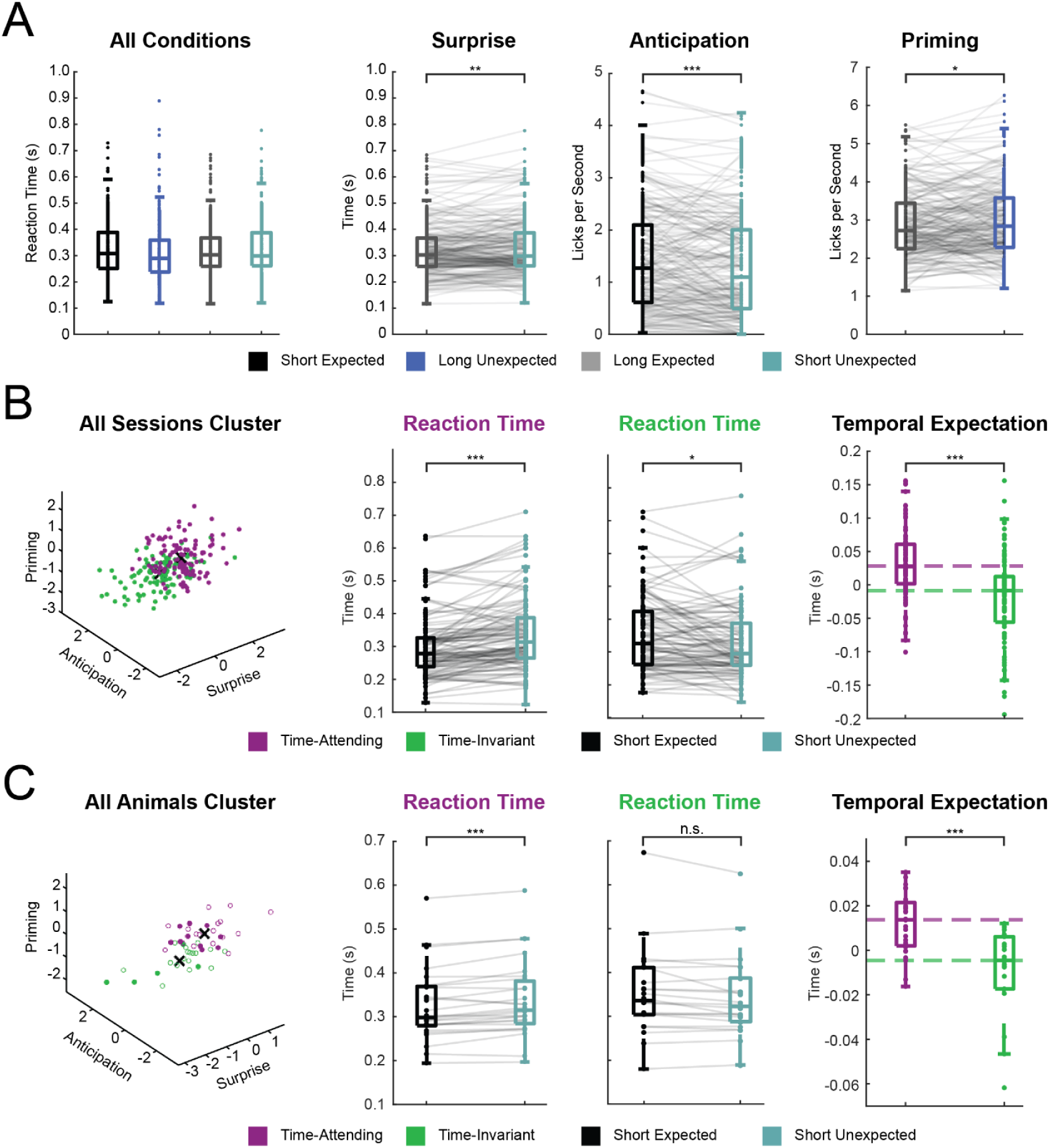
Clustering sessions and animals into time-attending and time-invariant groups. **(A)** Average reaction times per session for all four trial conditions. ‘Surprise’ measure based on reaction time comparing short unexpected and long expected trials (difference: 10 ms, p = 0.001, Wilcoxon sign rank, n = 216 sessions). ‘Anticipation’ measure based on licks per second in the cue-to-target interval [CTI] window comparing short unexpected and short expected trials (difference: 0.12 licks per second, p = 0.0002, Wilcoxon sign rank, n = 216 sessions). ‘Priming’ measure based on lick rate (lick events / s) in the pre-reward window comparing long unexpected and long expected trials (difference: 0.1 licks per second, p = 0.02, Wilcoxon sign rank, n = 216 sessions). **(B)** Clustering and classification of sessions according to the three temporal expectation measures described in (A): surprise, anticipation, and priming. Purple: time-attending cluster, green: time-invariant cluster. Average reaction times are shown, per session and across clustered groups (time-attending: p < 0.0001, Wilcoxon sign rank, n = 112 sessions; time-invariant: p = 0.014, Wilcoxon sign rank, n = 104 sessions), for conditions used to calculate TE-value: short unexpected trials - short expected trials. Comparison of average TE-values per session (p < 0.0001, Wilcoxon rank sum, n = 112 time-attending and 104 time-invariant sessions). **(C)** Clustering and classification of mice according to the three temporal expectation measures described in (A): surprise, anticipation, and priming. Purple: time-attending cluster, green: time-invariant cluster. Average reaction times are shown, averaged across sessions for each mouse and across clustered groups (time-attending: p = 0.0004, Wilcoxon sign rank, n = 22 animals; time-invariant: p = 0.094, Wilcoxon sign rank, n = 27 animals), for conditions used to calculate TE-value: short unexpected trials - short expected trials. Comparison of average TE-values per animal (p = 0.0001, Wilcoxon rank sum, n = 22 time-attending animals and 27 time-invariant animals).

### Stimulus-evoked fMRI BOLD responses vary with temporal attention before & after training

Regional dynamics across the time-attending and time-invariant groups was performed as above (see Fig. 3) by using a general linear model approach to assess whole-brain patterns of BOLD signal during se-fMRI before and after training. At baseline, we again find significant differences across groups, with generally higher positive BOLD contrast in the mice with time-invariant responses (Fig. 6A,C). When comparing these findings to the groups separated along the dimension of behavioural strategy (target-attending versus target-invariant, Fig. 3A,C), we find markedly similar baseline patterns of activation for both sets of analysis in response to the auditory cue stimulation, but the patterns of activation in response to the visual target stimulation and combined multisensory stimulation are markedly different in comparison to the behavioural strategies groups. Here, we find that the difference maps are generally less pronounced at baseline between the time-attending and time-invariant groups (Fig 6A). The significant differences (regionally individual t-values based on a p-value < 0.001 are listed in Suppl. Tabs. 4-5) that remain were largely in hippocampal-associated regions as well as in sensory (t-value = -5.12) and motor cortex (t-value = -5.93) (Fig. 6C).

**Figure 6.**
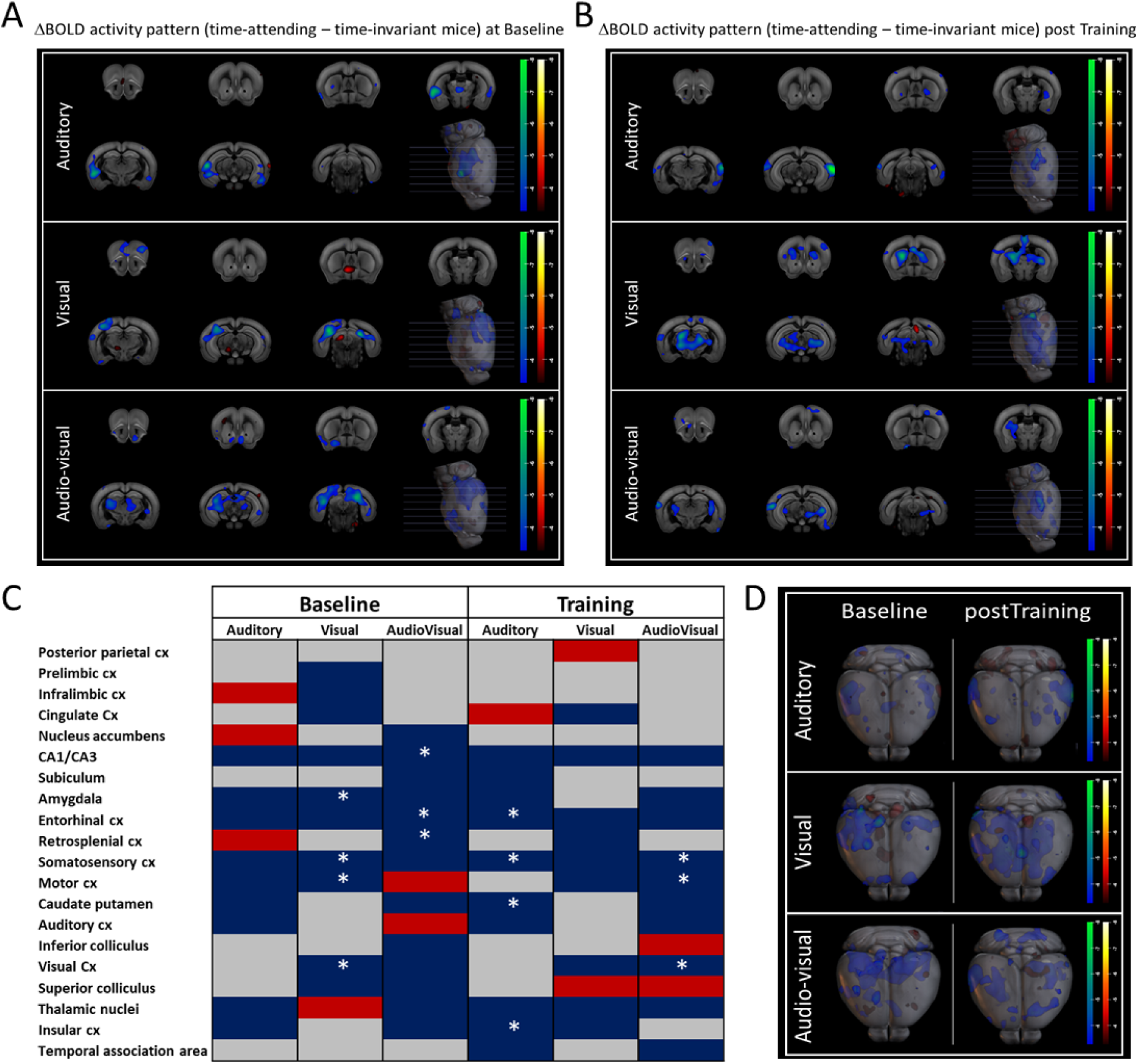
Quantification of whole-brain BOLD activity patterns for time-attending and time-invariant mice at baseline and post training. Difference maps of BOLD activity patterns for all three stimulation paradigms (auditory [cue], visual [target], audio-visual [simultaneous cue+target]) for mice with time-attending (n = 8) responses minus mice with time-invariant (n = 5) responses at baseline **(A)** and for mice with time-attending (n = 10) responses minus mice with time-invariant (n = 5) responses post training **(B)**. The significance level for the individual maps was set to p < 0.001 (t-value of 3.29) before analysis. The heatmap is based on the resultant t-values before subtraction of the images, here brighter colours indicate higher differences between the minuend and subtrahend map. The blue-green colour scale represents negative differences and the red-yellow colour scale positive differences between the BOLD contrasts of groups (time-attending - time-inattentive). **(C)** Observed differences of brain areas with detected differences for all three stimulation paradigms individually with their corresponding appearance (blue for negative difference [indicating that time-inattentive mice had higher BOLD response], red for positive difference [indicating that time-attending mice had higher BOLD response], grey marks undetectable regions). An asterisk highlights regions with statistical significant differences based on a two-sample unpaired t-test (p < 0.001). Individual t-values are listed in Supplementary Tables 4 and 5. **(D)** 3D volume rendering highlights the whole-brain differences with the same colour bars as shown in (A).

After training, while the time-invariant group continues to show higher levels of positive BOLD response, the patterns of activation between the two groups show significant changes (Fig. 6B-D). In particular, in response to the auditory cue stimulation, which shows significant differences in activation across various association cortices as well as the striatum (CP with t-value = -4.67) (Fig. 6C); this is in contrast to the behavioural strategies groups, which differed only in their activation of primary auditory cortex in response to the auditory cue after training (Fig. 3C). Suggesting that the auditory cue, which facilitates time-keeping for expectation-based behaviour during the task, plays a more prominent role to shape network plasticity in animals that attend to the temporal structure of the task. One of the few cortical regions that shows increased BOLD contrast in the time-attending group in response to the visual target was in posterior parietal regions, specifically the anterior secondary visual areas and bordering the posterior parietal association area (PPC; Fig. 6C), an association brain region (Lyamzin & Benucci, 2019; Whitlock, 2017) known to also play a role in attention (Malhotra et al., 2009), which may indicate the presence of unique neurophysiological activity in this cortical region in relation to temporal attention.

### Functional connectivity patterns at rest across mice with varying temporal attention

In relation to functional connectivity across the time-attending and time-invariant groups, we again investigated the activation during rs-fMRI for baseline and post training time points. In line with our se-fMRI results, the difference in overall correlation at baseline for these two temporal processing groups was lower (Fig. 7A,B; for mean correlation values ± se.e.m. see Suppl. Fig. 5) in comparison to the differences we observed at baseline for the groups that differed in their behavioural strategy (Fig. 4A,B). However, similar to the target-attending group, after training we found increased global correlations in time-attending mice (Fig. 7A,B) with a mean correlation value ± s.e.m. at baseline = 0.025 ± 5.7E-5 and post training = 0.043 ± 6.3E-5. Applying the same subnetwork graph theory approach to the time-attending versus time-invariant mice we found substantial differences in both the baseline as well as post training timepoints (Fig. 7C), and in comparison to the behavioural strategies groups (Fig. 4C). At baseline, a few regions show a similar node degree and amount of inter-regional connections, e.g., RSP, ACA and VIS for both time-attending and time-invariant groups; however, substantial differences in hippocampal-associated regions existed prior to training in these groups (Fig. 7C). After training, time attending mice showed increased functional connectivity within this subnetwork as evidenced by a higher number of edges and generally increased node degree, while time-invariant mice showed decreased functional connectivity after training as evidenced by fewer edges and generally decreased node degree. Interestingly, hippocampal-associated, cingulate cortex, and posterior parietal cortex nodes were differentially, altered in both groups after training, with time-attending mice showing a strengthening along these nodes (HPF = 20.20 ± 3.54; CG = 11.55 ± 3.58; PPC = 2.00 ± 1.51), and time-invariant mice showing a large decrease (HPF = 11.50 ± 3.64; CG = 5.30 ± 2.90; PPC = 0.50 ± 0.50) after training. Therefore, these results indicate an important neurophysiological link between frontoparietal and hippocampal circuits that relates directly to the availability of neural resources underlying task-related temporal attention.

**Figure 7.**
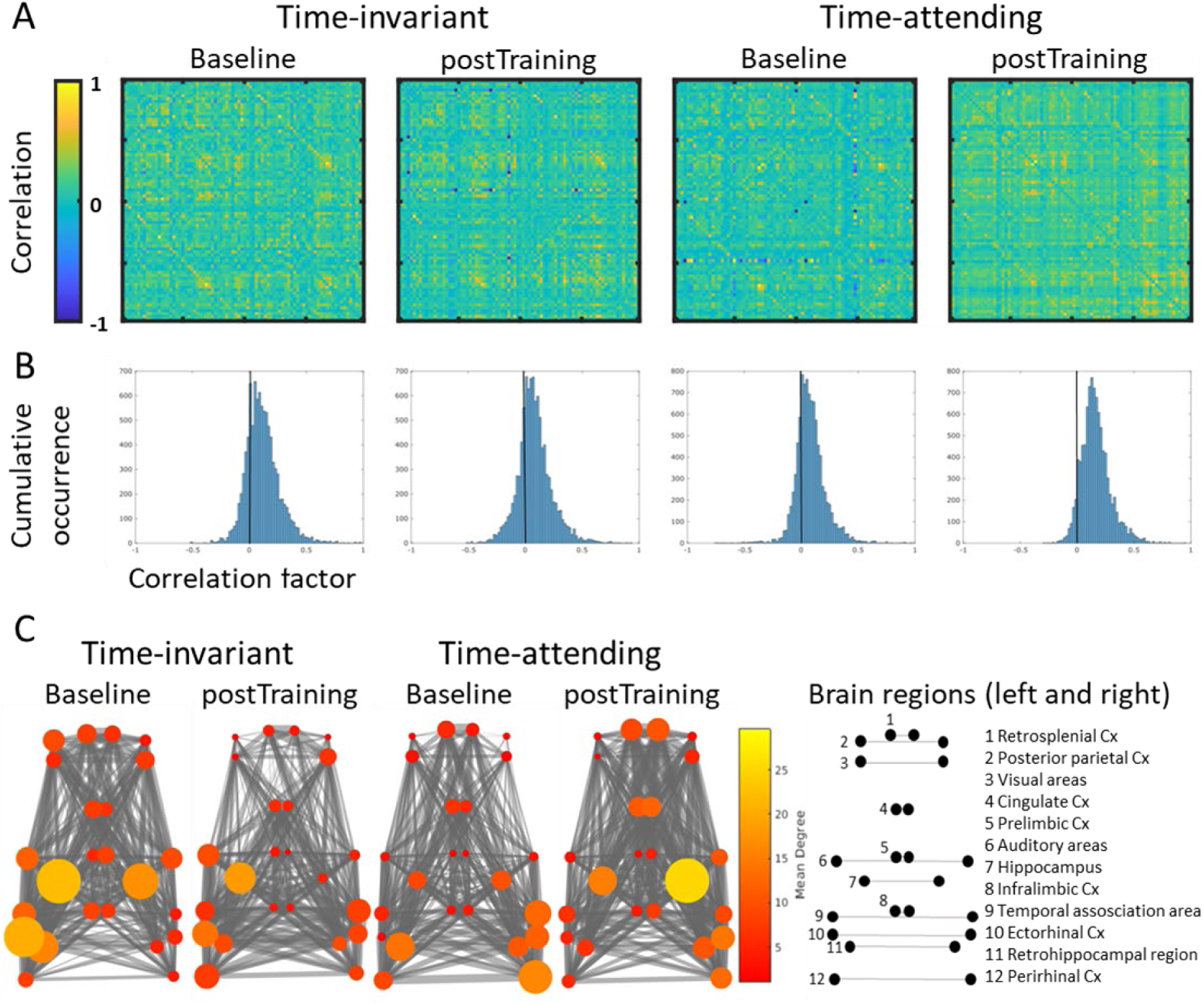
Functional connectivity of resting-state fMRI (rs-fMRI) before and after training for mice with time-attending versus time-invariant responses. **(A)** Correlation matrices across 49 brain regions bilaterally (98 total areas; see Suppl. Tab. 1) for time-invariant (n = 5) and time-attending (n = 10) mice calculated for baseline and post training rs-fMRI scans. Heatmap indicates the correlation strength based on the z-transform of the Pearson correlation coefficient (mean correlation value ± s.e.m. for time-invariant at baseline and post training = 0.035 ± 6.8E-5 and 0.029 ± 5.9E-5; mean correlation value ± s.e.m. for time-attending at baseline = 0.025 ± 5.7E-5 and post training = 0.043 ± 6.3E-5, see Suppl. Fig. 5). **(B)** Histogram of Pearson correlation coefficients across all 98 brain areas as calculated in (A). **(C)** A graph theory approach for selected brain areas of interest is shown across networks of interest (numbered list on right) for time-invariant and time-attending mice at baseline and post training rs-fMRI scans. Size and colour of the nodes represent the node degree (larger and brighter colour represents higher node degree). Mean node degree values ± s.e.m. of presented regions are available at Supplementary Figure 7.

### Representation of temporal expectation on the neuronal level

Finally, to further elucidate the neurophysiological basis of our functional connectivity findings, we investigated differences on the level of single-cell neuronal activity in the posterior parietal cortex (PPC) between time-attending and time-invariant mice. Two-photon calcium imaging was used to record single-cell neuronal activity in the PPC of mice. Chronic imaging of the same field-of-view (FOV) of neurons was performed during all testing sessions for mice that underwent cranial window implantation and GCaMP6m AAV injection (Fig. 8A, see methods). According to the same clustering approach described above (Fig. 5B), we classified each testing session that was performed with two-photon calcium imaging as time-attending or time-invariant (Fig. 8B; filled circles; n = 97 sessions, 49 time-attending, 48 time-invariant). We again confirmed that our clustering criteria recapitulated behavioural optimization of faster reaction times as defined by the TE-values; we found that TE-values for the time-attending sessions with calcium imaging were significantly higher than the corresponding time-invariant group (time-attending: 28 ± 7 ms; time-invariant: -28 ± 10 ms; p < 0.0001, Wilcoxon sign rank, n = 49 expecting and 48 time-invariant sessions; Fig. 8B). To evaluate whether neuronal activity within the PPC directly represents a readout of temporal expectation, we evaluated the decoding accuracy of the sensory-evoked neuronal activity that occurred following an expected versus an unexpected target stimulus, across both time-attending and time-invariant sessions (Fig. 8C). Indeed, template matching decoding (Montijn et al., 2014; van Duuren et al., 2008) resulted in a higher decoding accuracy between expected versus unexpected trials compared to randomly shuffled data across sessions for both time-attending (raw 60 ± 0.4 %; shuffled: 49 ± 0.03 %) as well as time-invariant (raw 58 ± 0.4 %; shuffled: 49 ± 0.03 %) sessions. Additionally, decoding accuracy in time-attending sessions was significantly higher than time-invariant sessions (p = 0.013, Wilcoxon sign rank, n = 49 expecting and 48 time-invariant sessions). Therefore, in line with our functional connectivity findings, neuronal activity within the PPC directly represents task-related temporal expectations, confirming the importance of this brain region within a wider functional network for temporal processing and temporal attention.

**Figure 8:**
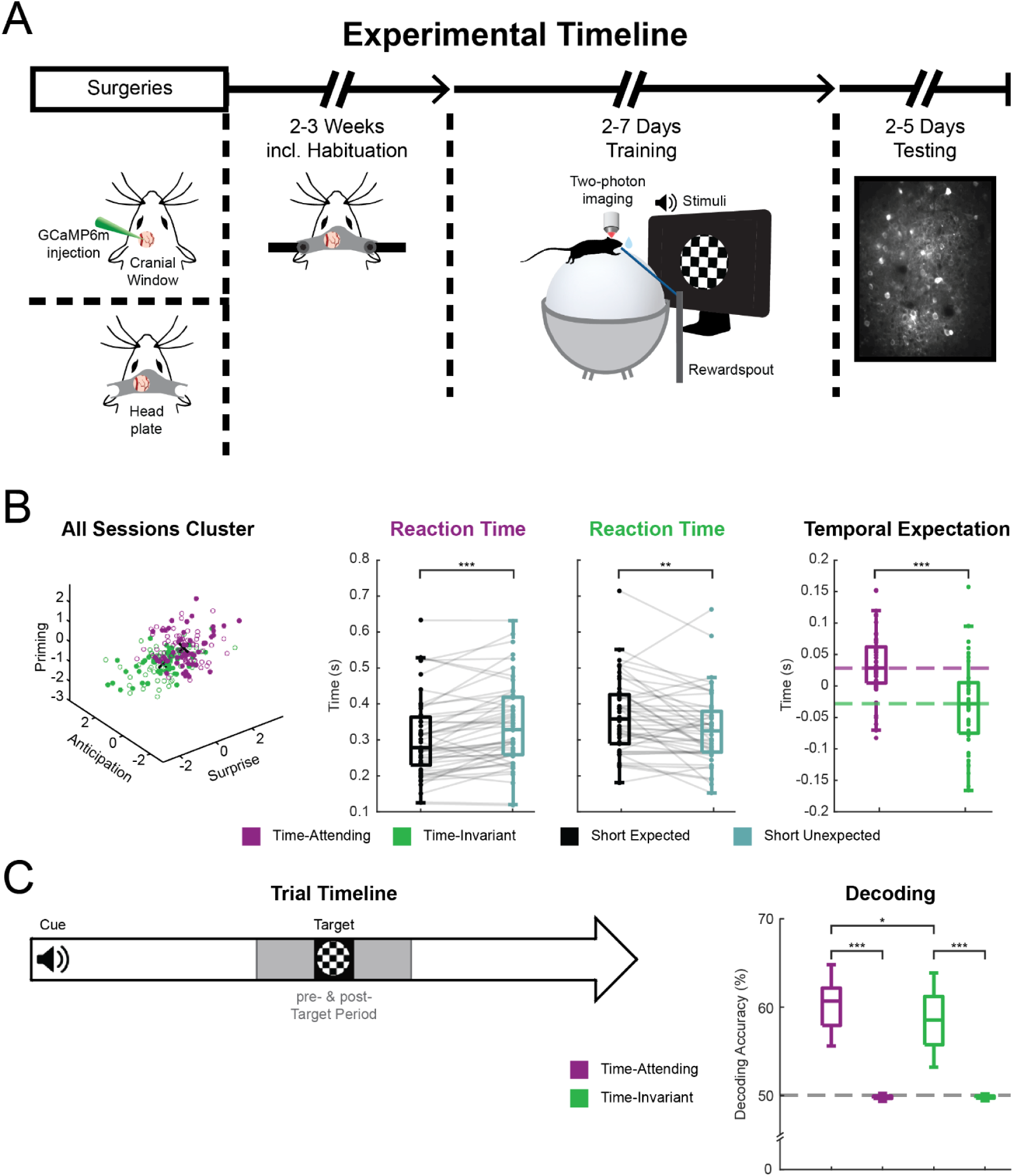
Neuronal activity in the posterior parietal cortex (PPC) on the single-cell level encodes for temporal expectation. **(A)** Experimental timeline including cranial window implantation to enable chronic two-photon calcium imaging during behaviour. **(B)** Clustering and classification of sessions according to the three temporal expectation measures: surprise, anticipation, and priming (see also Fig 5). Purple: time-attending cluster, green: time-invariant cluster. Average reaction times are shown, per session and across clustered groups (time-attending: p < 0.0001, Wilcoxon sign rank, n = 49 sessions; time-invariant: p = 0.002, Wilcoxon sign rank, n = 48 sessions), for conditions used to calculate TE-value: short unexpected trials - short expected trials. Comparison of average TE-values per session (p = 0.0001, Wilcoxon rank sum, n = 49 time-attending and 48 time-invariant sessions). **(C)** Schematic of pre-target, and post-target time windows during a trial. The difference in neural activity between time windows (post-pre) is the basis for the template-matching decoding approach. Decoding accuracies between expected versus unexpected trials for time-attending and time-invariant sessions is shown. Decoding was performed on raw data and shuffled datasets. Red horizontal line indicates the chance level of decoding accuracy. Decoding accuracies of raw datasets are significantly higher than their shuffled counterparts (time-attending: p < 0.0001, Wilcoxon sign rank, n = 49 sessions; time-invariant: p < 0.0001, Wilcoxon sign rank, n = 48 sessions) and decoding accuracy in time-attending sessions significantly higher than time-invariant sessions (p = 0.0137, Wilcoxon rank sum, n = 49 time-attending and 48 time-invariant sessions).

## Discussion

To optimize our actions, we must be able to predict when future events will occur. Here, we identified key networks that are differentially engaged during sensory-driven temporal processing depending on the adoption of particular behavioural strategies during learning, and further delineate how these networks support the formation of temporal expectations in order to optimize motor responses in mice. We established an experimental paradigm in which an expectation about the timing of sensory events is developed through statistical learning in mice. In humans, previous studies have used similar designs to show that an expected temporal delay results in a faster reaction time to a target stimulus because it occurs at an attended moment in time (Ball et al., 2022; Ball, Fuehrmann, et al., 2018; Ball, Michels, et al., 2018; Stefanics et al., 2010). We confirmed these results and found that, in mice, the processing of a stimulus presented at an expected moment in time is also facilitated, as readout by a decrease in reaction time as well as changes in anticipatory licking frequency. Additionally, the formation of temporal expectations relying on accumulated sensory information and multisensory cues involved integration via large macro-scale cortical networks comprised of primary sensory systems, association areas including posterior parietal cortex and retrosplenial cortex, prefrontal regions, as well as hippocampal networks.

Although mice showed evidence of attending to the temporal structure of the task, they also adopted divergent behavioural strategies and showed large interindividual variability in their behaviour. More efficient motor behaviours of confining licking to temporal periods directly related to the presentation of the visual target also resulted in the use of fewer sensory-evoked neural resources during fMRI both before and following task training. Generally, we found mainly positive BOLD responses for the target-invariant mice, even after training, which indicates a higher demand of neural resources to process the temporal structure of sensory stimuli. In contrast, target-attending mice showed less BOLD activity or even negative BOLD responses suggesting that they recruit less neural resources. While positive hemodynamic responses are generally accepted to reflect increased neuronal activity (Logothetis et al., 2001; A. J. Smith et al., 2002), negative responses are commonly observed but explanations of their origins are more controversial, e.g., whether they are associated with a suppression of neuronal activity (Bentley et al., 2016; Shmuel et al., 2006; Stefanovic et al., 2004) or more complex hemodynamics of the neurovascular system (He et al., 2022; Hu & Huang, 2015). There may also be regional differences since increases in hemodynamic and neuronal activity are both associated with positive BOLD signals in the cortex, but with negative BOLD signals in the rat hippocampus (Schridde et al., 2008). Negative bold signals have also been suggested to carry stimulus-specific information in visual cortex (Bressler et al., 2007), and shifts from positive to negative BOLD responses were also reported in rat superior colliculus (SC) upon varying discrimination performance of visual stimuli at different flickering frequencies (Gil et al., 2024). Thus, we conclude that positive and negative BOLD signals seen here in our target-invariant and target-attending mice, may reflect a respectively higher versus lower demand of neural resources when accomplishing the sensory-driven task.

Our findings of differential activation across time-invariant and time-attending mice within association cortices in posterior parietal regions, both on the functional network level and encoding on the single-cell level, as well as in the retrosplenial cortex, suggests that attending to this temporal structure of the task also recruited memory-guided attentional mechanisms (Lyamzin & Benucci, 2019; Malhotra et al., 2009; Whitlock, 2017). Here, we also found a role for frontoparietal networks, with associated changes in cingulate cortex, and increased functional connectivity and higher node degree specific to task-attentive mice between cingulate, posterior parietal and posterior-midline cortical regions. The involvement of posterior-midline association regions is also in line with recent descriptions of time cells in the retrosplenial cortex in rats (Subramanian & Smith, 2024), as well as findings that neurons within the retrosplenial cortex in mice have multidimensional representations of task structure that established a specific contextual encoding across learning (Sun et al., 2021).

The prominent role of hippocampal-associated networks was evident in their divergence across groups of mice that adopted different behavioural strategies during learning as well as in groups of mice that showed evidence of more or less reliance on the formation of temporal expectations to optimize motor behaviour. This is in line with previously described functions of the hippocampus in temporal processing as well as related sensory-driven representations (De Corte et al., 2022; Dylda & Pakan, 2021; Finnie et al., 2021; Pastalkova et al., 2008; Taxidis et al., 2020; Terada et al., 2017). Interestingly, here we show that pre-existing differences in BOLD responses and functional connectivity in hippocampal networks are strongly related to a later capacity for attending to the temporal structure of the task. Furthermore, these dynamics are differentially altered in hippocampal-associated networks over the course of training and in relation to both the capacity for attending to the temporal structure of the task as well as efficient learning strategies. In light of our results and the well-established importance of these hippocampal-associated networks in processes of aging and neurodegeneration (Adams et al., 2021; Billette et al., 2022; Caselli et al., 2009; Diersch et al., 2021; El Haj & Kapogiannis, 2016), future studies examining the link between sensory-driven temporal processing and the capacity for cognitive reserve in these neural circuits will also be of high interest.

## Materials and Methods

### Animals

Experiments were conducted on a total of 49 mice (24 males and 25 females, 2-6 months; 41 C57BL/6J wildtype mice (RRID: IMSR_JAX:000664; Jackson Laboratory), 8 Ai9 tdTomato reporter mice ([B6.Cg-Gt(ROSA)26Sortm9(CAG−tdTomato)Hze/J; RRID: IMSR_JAX:007909]). Mice were housed in standard cages with 12 h/12 h light dark cycle. All experiments were performed according to the NIH Guide for the Care and Use of Laboratory animals (2011) and the Directive of the European Communities Parliament and Council on the protection of animals used for scientific purposes (2010/63/EU) and were approved by the animal care committee of Sachsen-Anhalt, Germany (42502-2-1619 DZNE).

### Surgeries

Mice were anaesthetized with isoflurane (4% induction, ∼2% maintenance during surgery). Eye cream was applied to protect the eyes (Bepanthen, Bayer) and carprofen (5mg/kg) and dexamethasone (2 mg/kg) was injected subcutaneously. The scalp of the mouse was shaved and disinfected. The mouse was placed in a stereotaxic frame (Stoelting, USA) and body temperature was stabilized with a heating pad set to 37°C. If animals were used only for behavioural experiments, no craniotomy or adeno-associated virus (AAV) injections were performed, but the skull was exposed to attach a headplate to enable head-fixation.

### Cranial window and calcium indicator injection

After the skull was exposed, a craniotomy of 3 mm in diameter was made over the posterior parietal cortex (PPC, n = 28 mice). For calcium imaging an AAV was injected (AAV1.Syn.GCaMP6m.WPRE.SV40; RRID: Addgene_100841, titre: 2*10^12^) into the PPC (from Bregma: AP: -2.0 ML: 2.0). Injections were made with a Nanoject (World Precision Instruments, Germany) and a glass pipette with ∼20 μm tip diameter throughout the cortex in steps ranging from 200 µm to 700 μm in depth from the cortical surface. A total of 300 nl was injected. To prevent tissue damage, injection speed was set to 10 nl/min, and the pipette was kept in place for 1-5 minutes after each injection step. A glass coverslip was then placed over the craniotomy and fixed to the skull with glue (gel control, Loctite). A custom-built headplate was attached to the skull with glue and cemented with dental acrylic (Paladur, Heraeus Kulzer). Following the surgery, the animal was returned to its home cage for recovery. Behavioural training began 2-3 weeks following surgery to allow for virus expression and cranial window clearing (Dylda et al., 2018; Holtmaat et al., 2009).

### Behaviour

Before behavioural training, mice were handled for 7-10 days and habituated to the experimental setup and head-fixation for up to three days. During behavioural experiments, mice were water-restricted or given citric acid water (5%, (Urai et al., 2021); weight-loss was maintained under 15%) beginning 2-3 days before training and ending after the last recording. Mice received additional water based on sugar water rewards during the task. During training for 2-7 days mice were trained to lick a reward spout (200 µl of sugar water dispensed per trial) following the onset of a visual target stimulus (300 ms duration), which was presented 1.8 s after an auditory cue (8 kHz tone, 300 ms duration, ∼65 dB) for 200 trials per session. Trials were separated by an inter-trial-interval (ITI; 3 s). Mice were rewarded if they licked the reward spout within a time window (2 s) after target presentation. If mice licked before target presentation (i.e., during the CTI), a punishment in the form of a time-out (absence of stimuli, 3 s) was given before the start of the next trial. If licking occurred in the time-out period, the time-out was repeated until licking stopped for 3 s and the next trial was initiated. A trial only counted towards the 200 total trials, if at least the cue occurred. Not all animals received punishment during training. The last training day was followed by 2-5 testing days. Here, 200 trials in total were divided into two blocks of 100 trials. During testing, a short trial type with a CTI of 0.8 s was added to the training trial type with a long CTI of 1.8 s. Short and long CTI delay trials both blocks but with switched probabilities (either 75% or 25% probability), the order of which was pseudo-randomized. Block A had a 75% chance of long delay trials and 25% chance of short delay trials; Block B had a 25% chance of long delay trials and 75% chance of short delay trials. Block order was switched every day from the previous day starting with block A then block B on the first day.

During experiments, mice were situated on a 20 cm air-suspended polystyrene ball surrounded by six screens (JetBall-TFT, PhenoSys GmbH, Germany) covering 270° of the visual field. Loudspeakers (Genelec, Finland) were placed on both sides. Trials were conducted in the dark except when the specific target visual stimulus was presented. Movement was recorded using a JetBall system (PhenoSys GmbH, Germany). Licks were detected with a capacitive touch lick sensor (SEN-12041; Sparkfun, CO, USA) and recorded along with rewards, ball movements, and the onsets of sensory stimulation using PhenoSoft software (PhenoSys GmbH, Germany). A side-view camera (Imaging Source) captured the animal movements at 30Hz. PhenoSoft (PhenoSys GmbH, Germany) was used to synchronise and trigger the sensory stimulation, video monitoring, and start/stop the image acquisition of two-photon calcium imaging (using ScanImage software, Vidrio Technologies/MBF Bioscience).

### Two-photon imaging

Two-photon calcium imaging was performed using an 8 kHz resonant scanning microscope (HyperScope, Scientifica) with an Ultrafast laser (InSight X3 Dual output laser; Spectra-Physics; < 120 fs pulse width, 80 MHz repetition rate) tuned to 940 nm. Images were acquired at 30 Hz (using a 16X objective 0.8 NA, Nikon; tilted 5-10°) with ScanImage software (Vidrio Technologies/MBF Bioscience). Chronic calcium imaging of layer 2/3 neurons was performed at cortical depths of 180-220 µm. For each animal a single field of view was determined and followed over all training and testing days. Calcium imaging, video monitoring, and behavioural datafiles (movement encoders, lick times and stimulus onsets) were aligned post-hoc. Extracted regions of interest from the calcium imaging and pose-estimations from the behavioural video tracking (DeepLabCut, (Mathis et al., 2018), see below) were matched to the behavioural datafile by downsampling and interpolating such that the aligned datasets had the same number of total frames (overall sampling frequency ∼30Hz).

### Magnetic resonance imaging (MRI)

#### Animal preparation for MRI

For measurements before training, mice were anaesthetized with isoflurane (2 % in O_2_:N_2_ 1:1) and an intravenous catheter (inner diameter of the tube of 0.23 - 0.33mm, Instech Laboratories, USA, and a cannula with the size of 0,3 x 13mm, Braun Sterican, B.Braun, Germany) filled with a 1:5 medetomidine/NaCl solution (1 mg/mL, Orion Corporation, Finland/ 0.9 %, B.Braun, Germany) was placed into the tail vein. Afterwards, animals were transferred to the MRI system (BioSpec 94/20 USR, Bruker Biospin GmbH & Co. KG, Germany; for further details see below). For measurements after training, headplates of mice were removed under 4% isoflurane induction and 2% maintenance and the intravenous catheter was placed as described above.

#### General scanning procedure

Anatomical and functional MRI data were acquired using a 9.4 T Bruker AVANCE III small-animal MRI scanner (20 cm bore) equipped with a BGA12S gradient system (gradient strength: 600 mT/m, slew rate: 4570 T/m/s). A 112/086 ^1^H volume resonator was used for transmitting and a ^1^H 3x1 mouse optogenetic surface array coil for signal receiving. All parameters, data acquisition and reconstruction was controlled by ParaVision 7 software (Bruker Biospin GmbH & Co. KG, Germany).

Animals were placed on a mouse cradle with a bite bar and nose tube allowing gentle head fixation and inhalation gas delivery (for further details see below). The nose tube was placed with a short distance to the snout of the mouse to allow mounting a black/white circular gabor grid (similar to the one used in the behavioural experiments) on the bite bar. The mouse cradle itself was dyed in black colour to support light reflection. A pneumatic pillow was placed underneath the animal’s body for respiratory monitoring and a thermistor probe was placed rectally for body temperature monitoring. Vital parameters of the mice (breathing rate 120± 60 bpm, body temperature 37°C) were observed via PC-SAM software (Small Animal Instruments Inc., USA) and kept constant during the entire MR scan via a MR-compatible monitoring system (Model 1030, Small Animal Instruments, Inc., NY, USA) and an MR-compatible air heater system (Small Animal Instruments) consisting of a heater and a fan module and a hose fixated on the mouse cradle delivering warm air.

#### Sensory stimulation

Visual stimulation was delivered via an optic fibre cable placed nearby the right eye of the animal illuminating the gabor grid. The blue light of 1 Hz frequency (500 ms on/off) was generated by a LASER (l = 473 nm, 10mW, Thorlabs, Inc., USA) and controlled with a shutter system (Model SR470 and SR475, SRS).

Auditory stimulation was applied via two in-house modified piezo speakers (L010, frequency range 2 - 60 kHz, sound pressure level max. 120 dB, Kemo electronics, Germany) connected to hollow silicon tubes with pliable earplug tips (Pillow Soft Silicone Putty Earplugs, Mack’s, USA), which were placed through the openings of the surface array coil into the right and left ear channel of the animal. The stimulus (8 kHz tone, 500 ms on/off, 100 dB SPL) was generated by a LabView protocol (Version 2016, National Instruments, USA) and amplified (901 Personal Stereo Amplifier, Kramer Tools GmbH, Switzerland); the sound pressure level was controlled before each measurement via a handheld sound level metre (Sauter SU 130, Sauter GmbH, Germany).

For audio-visual stimulation, the auditory and visual stimulus was applied simultaneously.

All stimulation paradigms were controlled by means of dedicated LabView protocols and were temporally aligned throughout the scanning sequence to the scanner trigger signal via a digital input/output card (BNC-2110, National Instruments, USA).

#### Anatomical MRI

After a localizer scan for optimal animal placement, including all necessary adjustments such as basic frequency, global shim, reference power and receiver gain, a B0 map shim was applied to correct for B0 field inhomogeneities. Then, a 2D T2-weighted TurboRARE sequence (echo time/repetition time = 19/2241 ms, matrix = 256 x 256, 24 slices, resolution = 0.1 x 0.1 mm in plane, 0.6 mm slice thickness, 4 averages, scan time = 4 minutes) was acquired to gain an anatomical reference scan. This was used to confirm brain and skull integrity of mice and later alignment of functional MRI data.

#### Functional MRI

To compensate for cerebrovascular effects of isoflurane (Sullender et al., 2022; Van Aken et al., 1986), initial anaesthesia was switched to a lower concentration of isoflurane (0.5 % in 100% O_2_) in combination with the sedative medetomidine (i.v., 2 mg/ kg body weight/ hr, *via* tail vein catheter). After 30 min of administration, medetomidine dosage was further increased to 4 mg/kg body weight/hr. Scanning procedure started after an additional 30 min (i.e., 60 min after start of medetomidine administration) in order to ensure stable cerebrovascular conditions over the whole scanning time.

All fMRI scans were performed with a 2D GE-EPI sequence (echo time/repetition time = 16/ 1000 ms, matrix = 75 x 50, 24 slices, resolution = 0.4 x 0.4 x 0.6 mm, 5 dummy scans) after a 5 min gradient warm-up with the same sequence (to stabilise gradient performance). For se-fMRI, the number of repetitions was set to 600, for resting-state fMRI to 720 leading to total scanning times of 10 and 12 min, respectively. Three different se paradigms were applied always in the same order in a block-designed fashion with 120 s baseline recording followed by a stimulation period of 10 s stimulus and 40 s rest with in total 10 repetitions. First, the unimodal visual stimulation was performed, then the unimodal auditory stimulation and last the simultaneous audio-visual stimulation (for details see above). The resting-state scan was acquired after the se-fMRI scans. All functional MRI scans were exported to DICOM format *via* the ParaVison 7 software (Bruker).

### Data Analysis

#### Stimulus-evoked fMRI analysis

Se-fMRI data were analysed with Brainvoyager 20.6 software (Brain Innovation B.V., The Netherlands). Standard preprocessing pipelines were applied including removal of the first ten volumes, slice-time correction, 3D motion correction (three regressors for translation, three for rotation), a high-pass filter and 3D spatial smoothing of two pixels. One fMRI dataset for auditory stimulation after training was excluded from further analysis due to high motion artefacts and one mouse did not receive a second post training fMRI scan; this results in n = 15 animals for baseline and n = 14 animals for auditory and n = 15 animals for visual and audio-visual stimulation after training fMRI scan. Afterwards all anatomical scans were manually coregistered in 3D to the Allen Mouse Brain Atlas (Q. Wang et al., 2020) and the resultant transformation matrix including translation, rotation and scaling was applied to the equivalent functional scans to transfer them from the native space to the atlas space. A general linear model (GLM) approach was used for statistical group analysis (p<0.001, FDR-correction was not applied to keep the most relevant clusters) for all three stimulation paradigms individually. Here a canonical double-gamma hemodynamic response function with a time-to-response-peak of 2 s and a time-to-undershoot-peak of 10 s (adapted from (Lambers et al., 2020)) including a detection of autocorrelation and refitting of the GLM was used; therefore, animals were divided into two groups based on the behavioural measures (target- and time-attentive/ inattentive) to calculate the BOLD activity changes based on the learning effect. Finally, the statistical maps were exported to NIfTI format, the image pixels were set to 0.7 matching the atlas resolution for a correct overlay in ImageJ 1.52h (https://imagej.net/ij/). MRIcroGL 4.6 (https://www.nitrc.org/projects/mricrogl) was used for adequate visualization.

#### Resting-state fMRI analysis

Resting-state functional MRI scans were preprocessed with AIDAmri 2.0 (Pallast et al., 2019), which is a toolbox dedicated to mouse brains using adapted FSL 6.0 (Jenkinson et al., 2012; S. M. Smith et al., 2004; Woolrich et al., 2009) algorithms. Here, datasets underwent a bias correction (FSL FAST, (Zhang et al., 2001), brain extraction (FSL BET, (S. M. Smith, 2002), 3D motion correction (FSL FLIRT, (Jenkinson et al., 2002; Jenkinson & Smith, 2001), slice-time correction and coregistration to a customised brain reference atlas with 49 brain regions split between hemispheres based on the Allen Mouse Brain Atlas (Q. Wang et al., 2020) resulting in a total of 49 brain regions per hemisphere (see Suppl. Tab. 1). Connectivity analysis using graph theory was conducted with AIDAconnect 1.1. (Scharwächter et al., 2022), which is based on FSLNets v0.6, Niftyreg (Modat et al., 2010, 2014; Ourselin et al., 2001; Rueckert et al., 1999), the BCT toolbox (Rubinov & Sporns, 2010) and MATLAB 2023a (MathWorks, USA). Mice were separated again into the same two groups at baseline/ after training (n = 7/ 6 for attentive and n = 7/ 7 for inattentive mice) defined by behavioural outcomes. First, group correlation matrices were calculated based on the z-transform of the Pearson coefficient. Second, functional connections of some predefined regions of interest (posterior parietal, retrosplenial, cingulate, prelimbic, infralimbic, auditory, visual cortex, hippocampus, retrohippocampal area, temporal association area, ectorhinal and perirhinal cortex) were extracted as nodes and their functional connections as edges and arranged in a graph-theoretical network analysis graph. Last, several connectivity measures were computed (I) assortativity, (II) overall connectivity, (III) characteristic path length and (IV) global efficiency. For illustration, all images were arranged in Adobe Illustrator 28.3 (Adobe Inc., USA).

#### Behaviour analysis

Several important time windows during trials were delineated by the trial structure: i) cue-to-target Interval (CTI): time between the onset of the auditory cue and the onset of the visual target; ii) reward window: 2 s time window starting after the onset of the target stimulus in which a lick event would trigger the delivery of a water reward; iii) intertrial interval (ITI): the time between the end of the reward window and the onset of the auditory cue of the following trial. Additional time windows were used for analysis: i) cued delay window: the time following the onset of the auditory cue until 250 ms before the target onset, in order to avoid effects of anticipatory licking; ii) pre-reward window: time period from 250 ms before the target onset until the reward is given, in order to capture dynamics related to anticipatory behaviour; iii) target windows: pre-target window is the time period from 500 ms before the target onset and post-target window is the 500 ms period following the target onset, used to normalize target-evoked analysis of both licking rate as well as calcium activity.

A reaction time for each trial was calculated as the time between the onset of the visual target stimulus to the first lick event following this. If no lick event occurred within the given reward window (2 s), the trial was considered a missed trial and was excluded from average reaction time measures. Lick rate measures were calculated by summing the total number of lick events that occurred in a given time window and dividing by the extent of the given time window in seconds (i.e., lick events / s).

All reported differences in both reaction time measures as well as lick rate measures between two trial conditions was calculated by taking the median of the population of responses per trial (Equation 1).

**Equation 1: Differences between conditions.** The difference of a measure between two conditions (e.g., short expected trials [*Condition1*] and short unexpected trials [*Condition2*]) is defined as the median of differences between *Condition1* and *Condition2* of all trial data points in a population *X*, as follows:

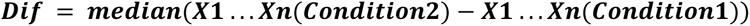

The target modulation index for the lick rate was calculated as the target-evoked normalized change in lick rate, as defined in Equation 2.

**Equation 2: Target modulation index of lick rate.** Defined as the normalized change in the licks per second (*LpS)* between the *pre*-target time window (LpS_pre_) and *post*-target reward window (LpS_post_). Pre-target window = 0.5 s before target onset until time of target, post-reward window = target onset until 0.5 s after target.

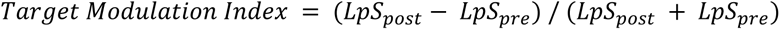

To classify individual daily behavioural sessions as target-attending versus target-invariant behavioural strategies, raw licking behaviour (such as that shown in Fig. 1B) was examined by two independent observers for all sessions to determine if the behaviour in the session showed spatially confined licking behaviour in relation to the target onset, or if the licking behaviour spanned the extent of the trial duration. In this way, factors such as missed trials, within session and across-trial dynamics, and changes in patterns of behaviour across blocks could also be taken into consideration in assigning a behavioural strategy to the session as a whole. Since individual mice could demonstrate either behavioural strategy on a given session and or change strategy across daily sessions, to classify individual mice as adopting an overall more target-attending versus an overall more target-invariant behavioural strategy, the average target modulation index across sessions was calculated for each mouse and a threshold of ≥ 0.3 was used to classify mice as target-attending as this represents an overall doubling in the target-evoked rate of licking. Mice with an average target modulation index of the lick rate of ≤ 0.3 across sessions were considered as adopting an overall target-invariant behavioural strategy.

#### Two-photon analysis

Images collected during two-photon calcium imaging were analysed as previously described (Henschke et al., 2020; Pakan, Currie, et al., 2018; Pakan et al., 2016). Briefly, images were motion corrected using discrete Fourier 2D-based image alignment of image frames (SIMA 1.3.2, sequential image analysis (Kaifosh et al., 2014)). Regions of interest (ROIs) were selected manually and corresponded to all visually identifiable neuronal cell bodies in images downsampled to 1 Hz. During ROI segmentation, experimenters are given generic animal IDs and blind to experimental timepoints within the data (i.e., stimulus onsets, behavioural state, etc.). The pixel intensity was then averaged within each ROI to create a raw fluorescence time series F(t). Baseline fluorescence F0 was computed for each neuron by taking the 5th percentile of the smoothed F (t) (1 Hz lowpass, zero-phase, 60th-order FIR filter) and the change in fluorescence relative to baseline (ΔF/F_0_) was calculated (F(t)-F0/F0). We used nonnegative matrix factorization (NMF), in order to remove neuropil contamination as implemented in FISSA (Keemink et al., 2018). All further analyses were performed using custom-written scripts in MATLAB 2022b (MathWorks, MA, USA).

Template matching decoding methods were used to calculate decoding accuracy between unexpected trials (Montijn et al., 2014; van Duuren et al., 2008). Target-evoked average ΔF/F_0_ was normalized using a 0.5 second time window before (pre) and after (post) each presented visual target stimulus (post-pre) per trial and this normalized ΔF/F_0_ was used for the template-matching decoding. Decoding was performed on raw data and shuffled datasets as follows: For each session an equal number of trials was picked from each condition (maximum number of trials in condition with fewest trials) at random. This process was performed 1000 times per session and the results were averaged. For the shuffled dataset all trials across both conditions were first pooled and then randomly assigned either condition before performing the same decoding process as described above.

#### Video analysis

Pose-estimations and frame-by-frame tracking were performed on the video monitoring that was captured during the behavioural task. The side-view of the body of the animal was analysed using DeepLabCut (DLC) (Mathis et al., 2018). In short, we trained DLC on the videos resulting from the side-view camera to track limbs, joints, tail and facial features (e.g., tongue, whiskers and jaw) of mice as well as the lick spout with individual markers. The DLC analysis was then used to determine frames of the videos where the animal was grooming using their forepaws (i.e., cleaning their face/whiskers). Grooming was defined as a point in time where the animal’s forepaw was within a 25 pixel radius of the lick spout. Since the lick spout is a capacitive sensor, the forepaw touching the lick spout will be recorded as a lick by the capacitive touch sensor. We removed grooming-related events from the dataset if grooming, detected by DLC, coincided with the same frame a lick event was recorded by the capacitive touch sensor.

### Statistics

Statistical tests used are indicated in the results and conducted in MATLAB. Statistical significance was assumed at p<0.05. One-, and two-way ANOVA were used to determine mixed effects of trial structure and/or groups on reported data. Unless otherwise specified, average values are reported as mean ± s.e.m.

## Acknowledgements

This work was supported by the Deutsche Forschungsgemeinschaft (DFG) Collaborative Research Center (CRC) 1436 - project 425899996 (subproject B06). We thank Dr. Michael Lippert for technical support in LabView programming as well as Janet Stallman, Anja Gürke and Cathleen Knape for technical assistance. We thank the GENIE Program and the Janelia Research Campus for making GCaMP6 available.

## Author Contributions

TN, JMPP and EB acquired funding for the project and conceived of the experiments, with additions to experimental design from JUH, AW and PW. Experiments were performed by AW and JH (behaviour and calcium imaging) and AM and PW (MRI experiments). Data was analyzed by AW and PW in consultation with JUH, JMPP and EB. Results and interpretation were done by AW, JUH, JMPP, PW and EB and discussed with all authors. AW, PW, JMPP and EB wrote the manuscript with input from all authors.

## Abbreviation list

2D: two-dimensional
3D: three-dimensional
AAV: adeno-associated virus
ACB: nucleus accumbens
ACA: anterior cingulate area
Amy: amygdala
ANT: anterior nucleus
APN: anterior pretectal nucleus
AUD: auditory areas
BOLD: blood-oxygen-level-dependent
bpm: breaths per minute
CA1: cornu ammonis 1
CA3: cornu ammonis 3
CG: cingulate area
cm: centimetre
CP: caudoputamen
CRC: Collaborative Research Centre
CTI: cue-target-interval
cx: cortex
°C: degree Celsius
DAPI: 4′,6-diamidino-2-phenylindole
dB: decibel
DFG: Deutsche Forschungsgemeinschaft
DICOM: digital imaging and communication
DLC: DeepLabCut
DREADD: designer receptors exclusively activated by designer drugs
ECT: ectorhinal cortex
ENT: entorhinal cortex
FDR: false discovery rate
Fig: figure
FIR: finite impulse response
fMRI: functional magnetic resonance imaging
fs: femtoseconds
GLM: general linear model
g: gramm
HPF: hippocampal formation
hr: hour
Hz: hertz
IC: inferior colliculi
ILA: infralimbic area
i.p.: intraperitoneally
ITI: inter-trial-interval
i.v.: intravenously
ID: identification number
kHz: kilohertz
kg: kilogram
LD: laterodorsal nucleus
LG: lateral geniculate complex
m: metre
MD: mediodorsal nucleus
mg: milligram
MHz: megahertz
min: minutes
mL: millilitre
mm: millimetre
MoCx: motor cortex
MR: magnetic resonance
MRI: magnetic resonance imaging
ms: milliseconds
mT: millitesla
mW: milliwatt
m: micrometre
NA: numerical aperture
NIfTI: neuroimaging informatics technology initiative
nL: nanoliter
nm: nanometre
NMF: nonnegative matrix factorization
PBS: phosphate-buffered solution
PERI: perirhinal cortex
PFA: paraformaldehyde
PFC: prefrontal cortex
PL: prelimbic area
PMT: photomultiplier tube
PO: posterior complex of the thalamus
PPC: posterior parietal cortex
PPT: posterior pretectal nucleus
RHP: retrohippocampal region
ROI: region of interest
rs: resting-state
RSC: retrosplenial cortex
RT: reticular nucleus
s: second
SC: superior colliculus
se: stimulus-evoked
s.e.m: standard error of the mean
SSCx: somatosensory cortex
SSp: primary somatosensory cortex
SSs: supplementary somatosensory cortex
SUB: subiculum
T: Tesla
TE: temporal expectation
TEa: temporal association area
VIS: visual areas
VT: ventral nucleus

## Supplemental Information

### Supplementary Tables

**Supplementary Table 1.**
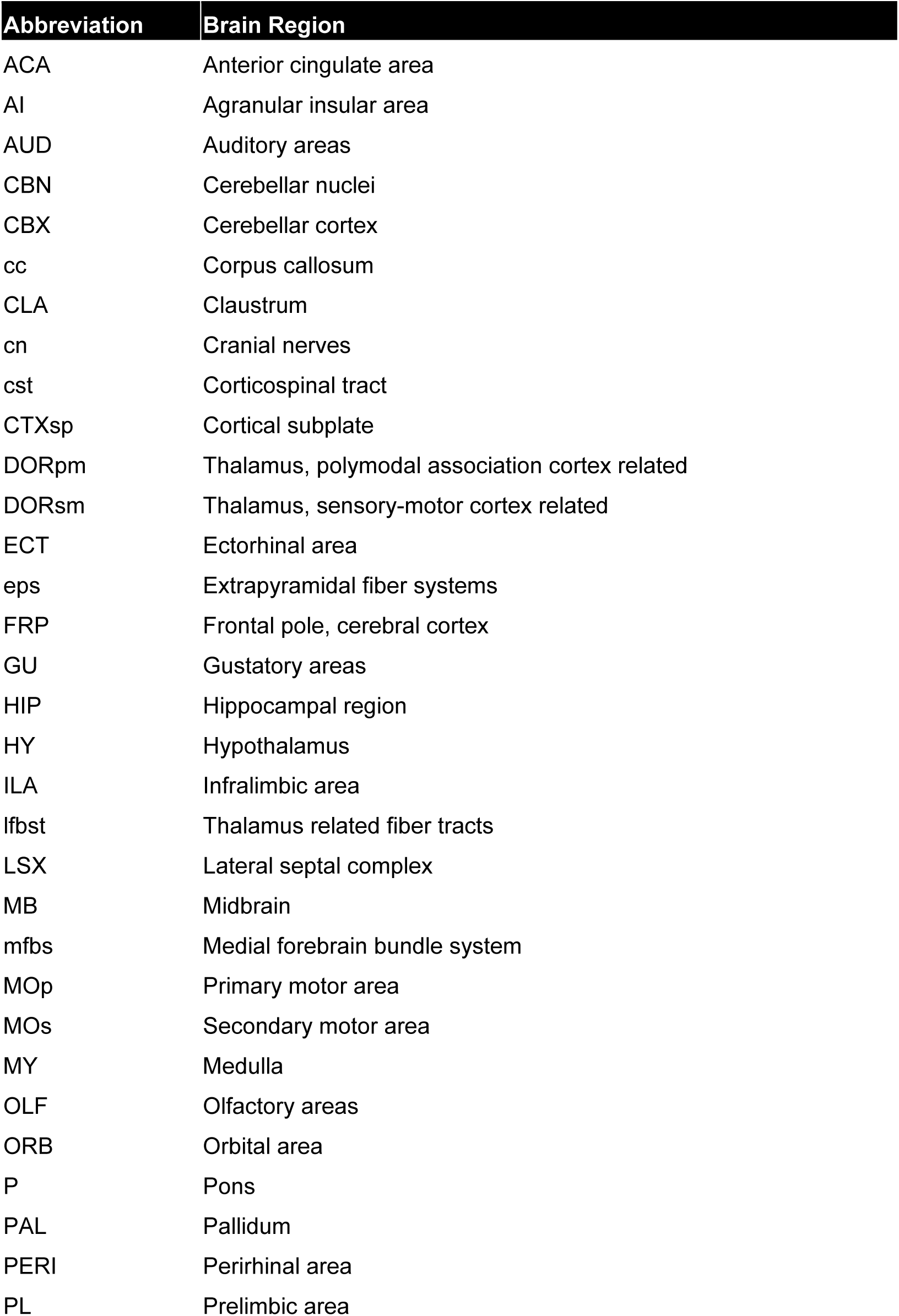
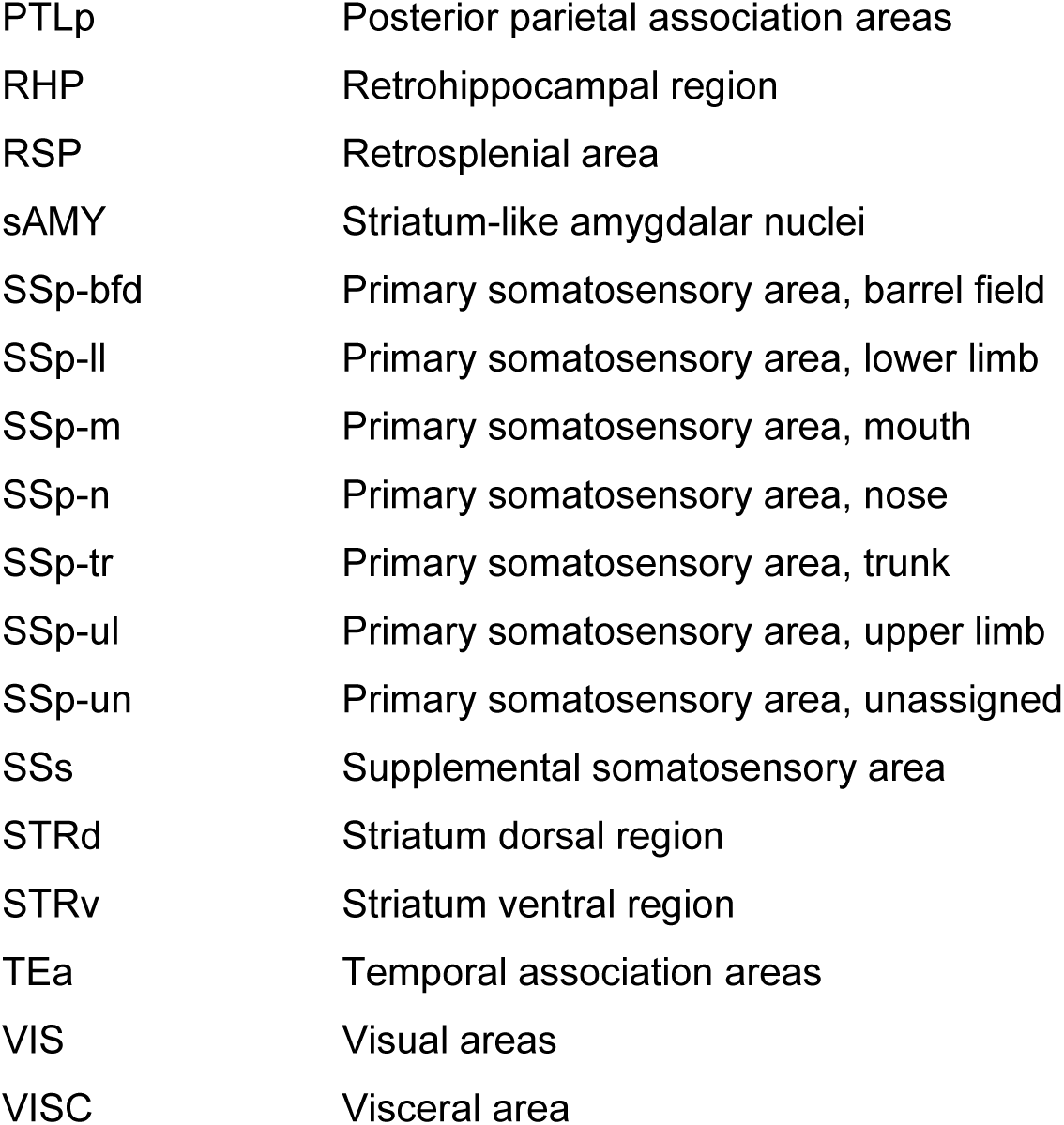
List of the 49 brain regions per hemisphere derived from the Allen Mouse Brain Atlas for resting-state fMRI analysis.

| Abbreviation | Brain Region |
| --- | --- |
| ACA | Anterior cingulate area |
| AI | Agranular insular area |
| AUD | Auditory areas |
| CBN | Cerebellar nuclei |
| CBX | Cerebellar cortex |
| cc | Corpus callosum |
| CLA | Clastrum |
| cn | Cranial nerves |
| cst | Corticospinal tract |
| CTXsp | Cortical subplate |
| DORpm | Thalamus, polymodal association cortex related |
| DORsm | Thalamus, sensory-motor cortex related |
| ECT | Ectorhinal area |
| eps | Extrapyramidal fiber systems |
| FRP | Frontal pole, cerebral cortex |
| GU | Gustatory areas |
| HIP | Hippocampal region |
| HY | Hypothalamus |
| ILA | Infralimbic area |
| lfbst | Thalamus related fiber tracts |
| LSX | Lateral septal complex |
| MB | Midbrain |
| mfbs | Medial forebrain bundle system |
| MOp | Primary motor area |
| MOs | Secondary motor area |
| MY | Medulla |
| OLF | Olfactory areas |
| ORB | Orbital area |
| P | Pons |
| PAL | Pallidum |
| PERI | Perirhinal area |
| PL | Prelimbic area |
| PTLp | Posterior parietal association areas |
| RHP | Retrohippocampal region |
| RSP | Retrosplenial area |
| sAMY | Striatum-like amygdalar nuclei |
| SSp-bfd | Primary somatosensory area, barrel field |
| SSp-lI | Primary somatosensory area, lower limb |
| SSp-m | Primary somatosensory area, mouth |
| SSp-n | Primary somatosensory area, nose |
| SSp-tr | Primary somatosensory area, trunk |
| SSp-ul | Primary somatosensory area, upper limb |
| SSp-un | Primary somatosensory area, unassigned |
| SSs | Supplemental somatosensory area |
| STRd | Striatum dorsal region |
| STRv | Striatum ventral region |
| TEa | Temporal association areas |
| VIS | Visual areas |
| VISC | Visceral area |

**Supplementary Table 2.**
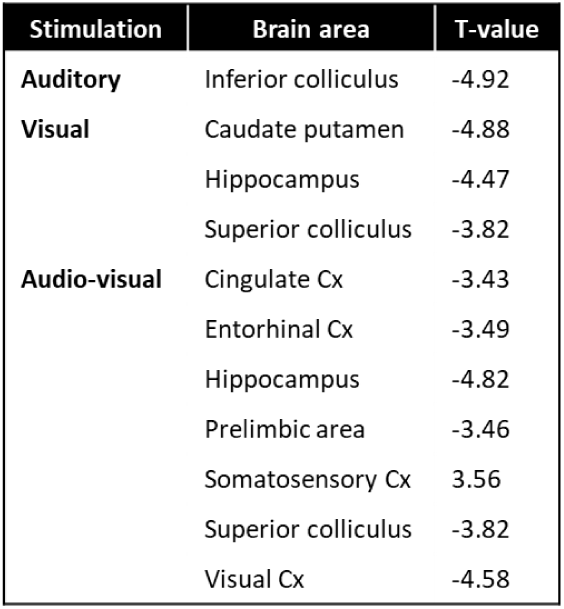
T-values for significantly different brain regions based on a two-sample t-test for the group results of se-fMRI for auditory, visual and audio-visual stimulation for target-attending – target-invariant mice at baseline. Negative values correspond to a significant negative difference.

**Supplementary Table 3.** T-values for significantly different brain regions based on a two-sample t-test for the group results of se-fMRI for auditory, visual and audio-visual stimulation for target-attending – target-invariant mice post training. Negative values correspond to a significant negative difference.

| Stimulation | Brain area | T-value |
| --- | --- | --- |
| Auditory | Auditory Cx | -4.92 |
| Visual | Caudate putamen | -4.88 |
|  | Hippocampus | -4.47 |
| Audio-visual | Amygdala | 3.81 |
|  | Auditory Cx | -6.33 |
|  | Caudate putamen | -6.57 |
|  | Cingulate Cx | -5.73 |
|  | Entorhinal Cx | -3.53 |
|  | Hippocampus | -6.64 |
|  | Motor Cx | -5.64 |
|  | Prelimbic area | -6.24 |
|  | Retrosplenial Cx | -3.45 |
|  | Somatosensory Cx | -4.70 |
|  | Superior colliculus | -6.49 |
|  | Visual Cx | -4.28 |

**Supplementary Table 4.** T-values for significantly different brain regions based on a two-sample t-test for the group results of se-fMRI for auditory, visual and audio-visual stimulation for time-attending – time-invariant mice at baseline. Negative values correspond to a significant negative difference. The x indicates no significant difference for this stimulation paradigm.

| Stimulation | Brain area | T-value |
| --- | --- | --- |
| **Auditory** | x | x |
| **Visual** | Amygdala | -4.91 |
|  | Motor Cx | -5.93 |
|  | Somatosensory Cx | -5.12 |
|  | Visual Cx | -5.22 |
| **Audio-visual** | CA1/CA3 | -4.64 |
|  | Entorhinal Cx | -5.28 |
|  | Retrosplenial Cx | -4.66 |

**Supplementary Table 5.** T-values for significantly different brain regions based on a two-sample t-test for the group results of se-fMRI for auditory, visual and audio-visual stimulation for time-attending – time-invariant mice post training. Negative values correspond to a significant negative difference. The x indicates no significant difference for this stimulation paradigm.

| Stimulation | Brain area | T-value |
| --- | --- | --- |
| **Auditory** | Caudate putamen | -4.67 |
|  | Entorhinal Cx | -4.72 |
|  | Inferior colliculus | -5.48 |
|  | Insular Cx | -5.20 |
|  | Somatosensory Cx | -4.45 |
| **Visual** | x | x |
| **Audio-visual** | Motor cx | -5.44 |
|  | Retrosplenial Cx | -6.69 |
|  | Somatosensory Cx | 5.04 |
|  | Visual Cx | -5.39 |

### Supplemental Figures

**Supplementary Figure 1.**
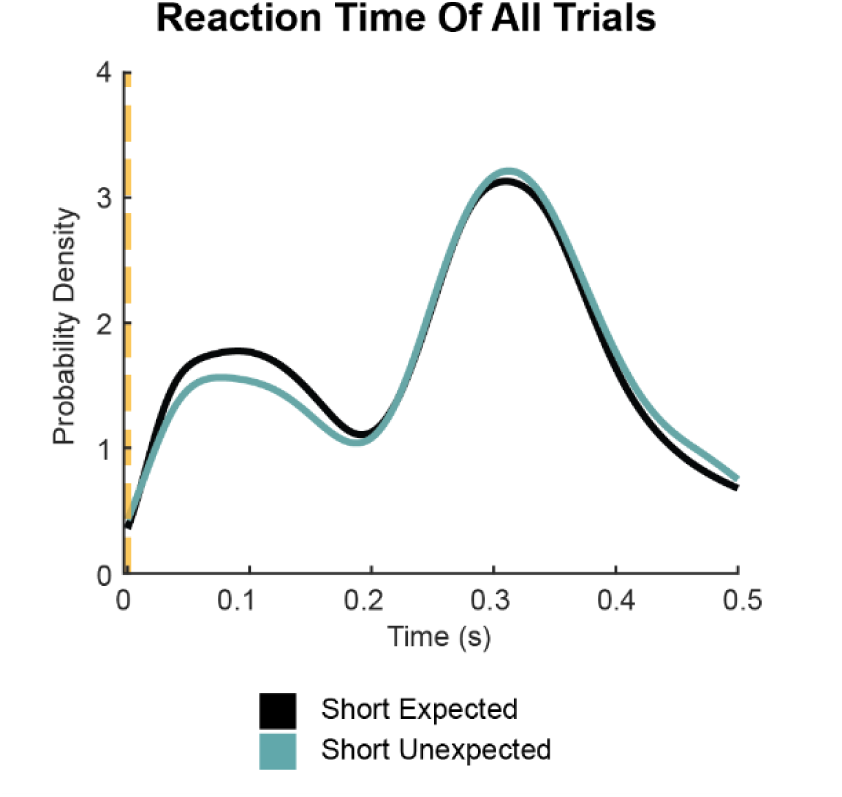
Probability density of all reaction times. Probability density distribution of reaction times for all trials in the dataset. Black: Short expected; teal: Short unexpected. Probability density of all reaction times for short expected and short unexpected trials in the dataset (n = 16,091 short expected and 5,417 short unexpected trials). Yellow dashed line, target onset.

**Supplementary Figure 2.**
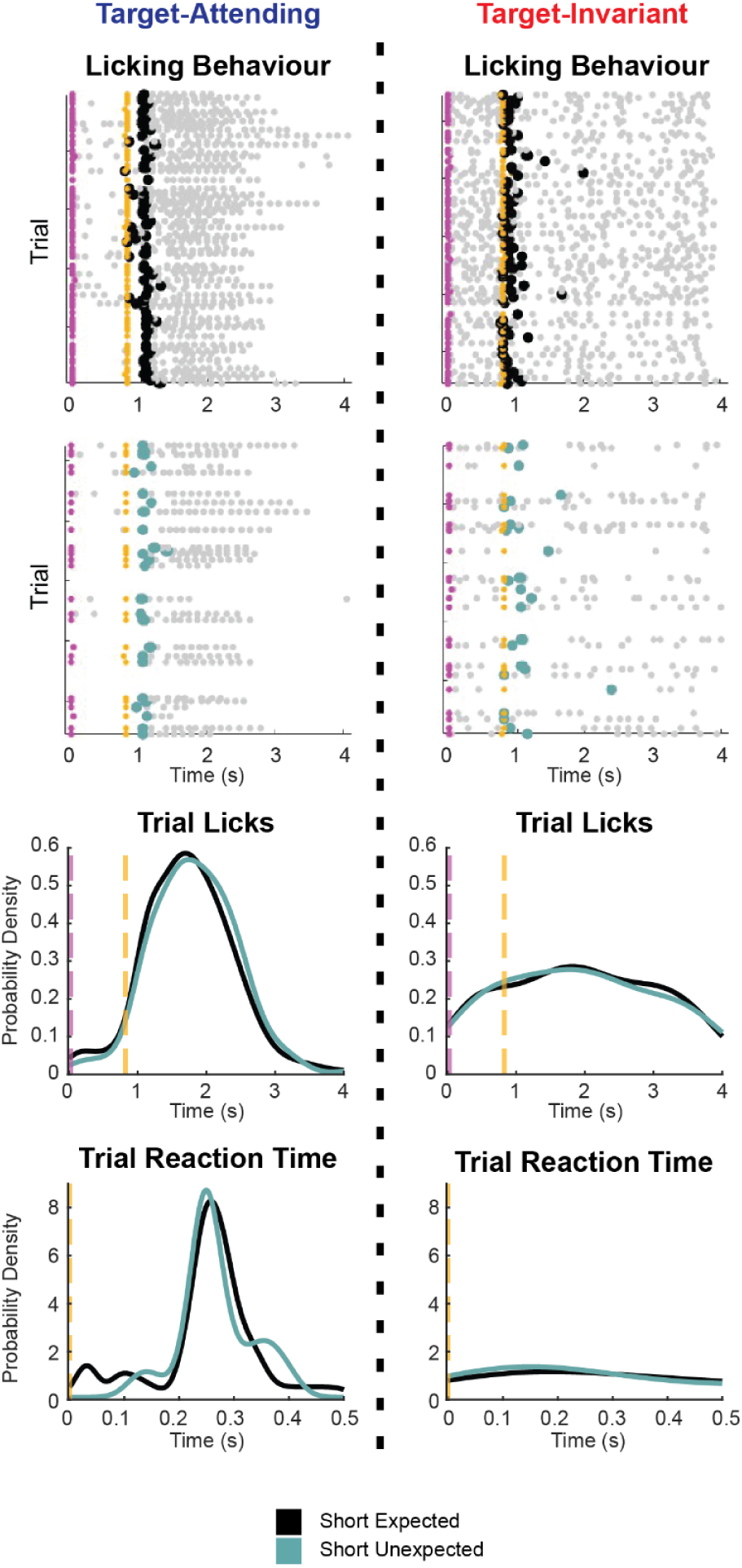
Example behaviour of target-attending licking and target-invariant licking. Two additional example sessions of target-attending and target-invariant licking behaviour during short delay trials across blocks. *Top*: raw licking events across blocks. Pink dots: cue onset per trial. Yellow dots: target onset per trial. Grey dots: lick events. The first lick occuring after the target (large coloured dots) was used to calculate reaction time per trial. *Bottom*: probability density of all lick events, and probability density of all reaction times per trial for the corresponding example session. No significant difference in short expected and short unexpected reaction times was found for both example sessions. Target-attending example: TE-value: < 0.001 ms, p = 0.62, Wilcoxon rank sum, n = 75 expected and n = 25 unexpected trials; target-invariant example: TE-value: -125 ms, p = 0.45, Wilcoxon rank sum, lick: n = 74 expected and n = 24 unexpected trials.

**Supplementary Figure 3.**
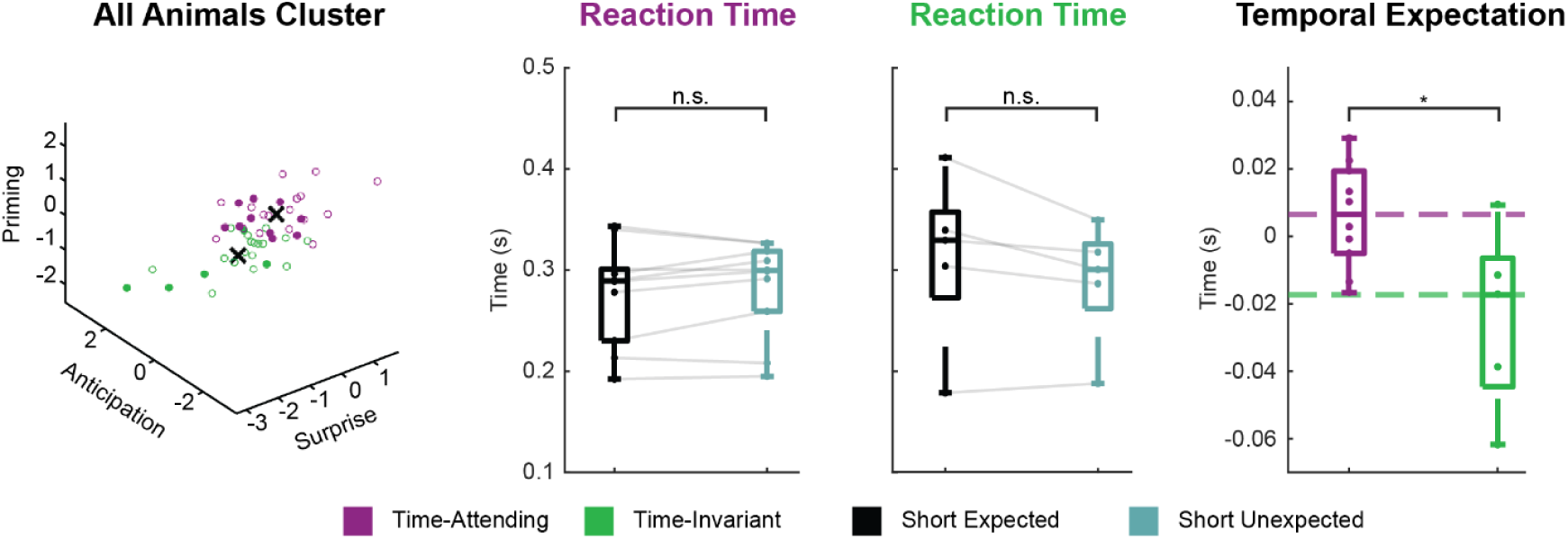
Evaluation time-attending and time-invariant TE-value of only MRI mice. Clustering and classification of sessions according to the three temporal expectation measures: surprise, anticipation, and priming (see also Fig 5). Purple: time-attending cluster, green: time-invariant cluster. Average reaction times are shown, averaged across sessions for each MRI mouse and across clustered groups (time-attending: p = 0.3, Wilcoxon sign rank, n = 10 animals; time-invariant: p = 0.1, Wilcoxon sign rank, n = 5 animals), for conditions used to calculate TE-value: short unexpected trials - short expected trials. Comparison of average TE-values per animal (p = 0.02, Wilcoxon rank sum, n = 10 time-attending animals and 5 time-invariant animals).

**Supplementary Figure 4.**
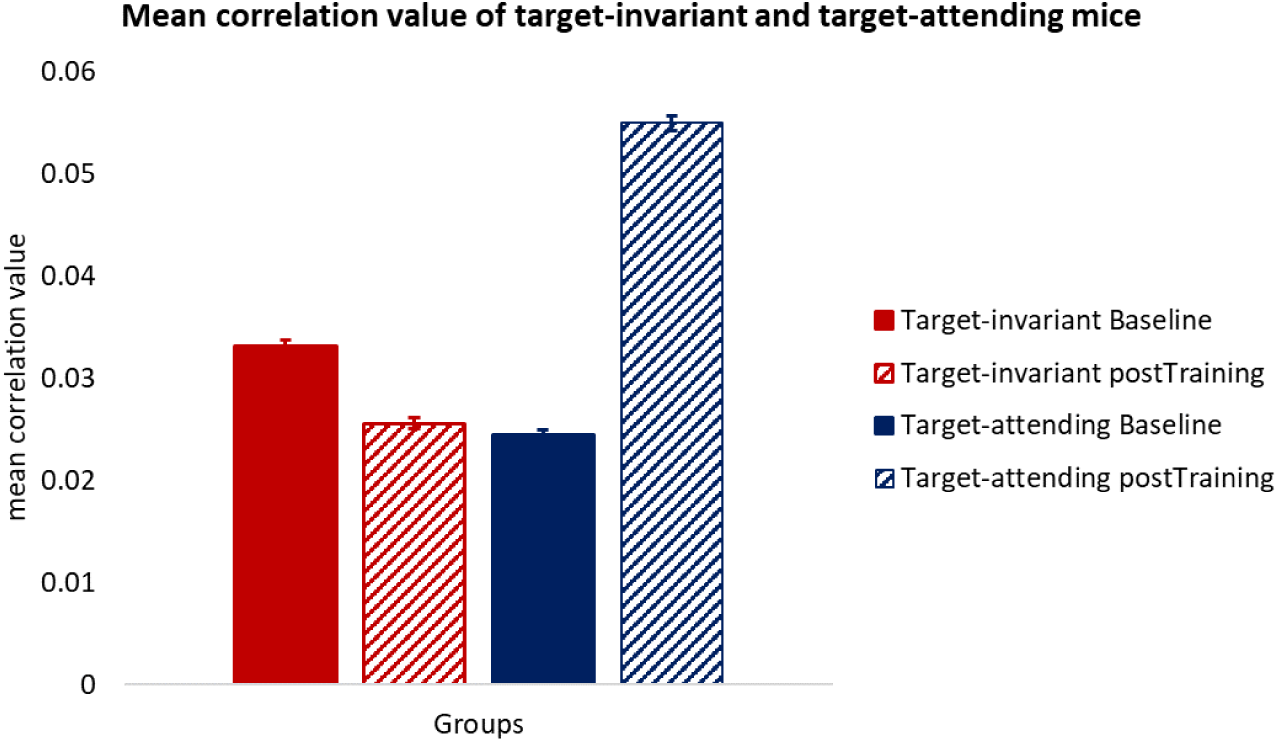
Mean correlation values of target-invariant and target-attending mice based on correlation analyses of the resting-state fMRI. The mean correlation coefficient and the corresponding s.e.m. were calculated for both groups of animals (target-invariant [red], target-attending [blue]) at baseline (filled) and post training (shaded) based on their correlation values extracted from the individual Pearson correlation matrix.

**Supplementary Figure 5.**
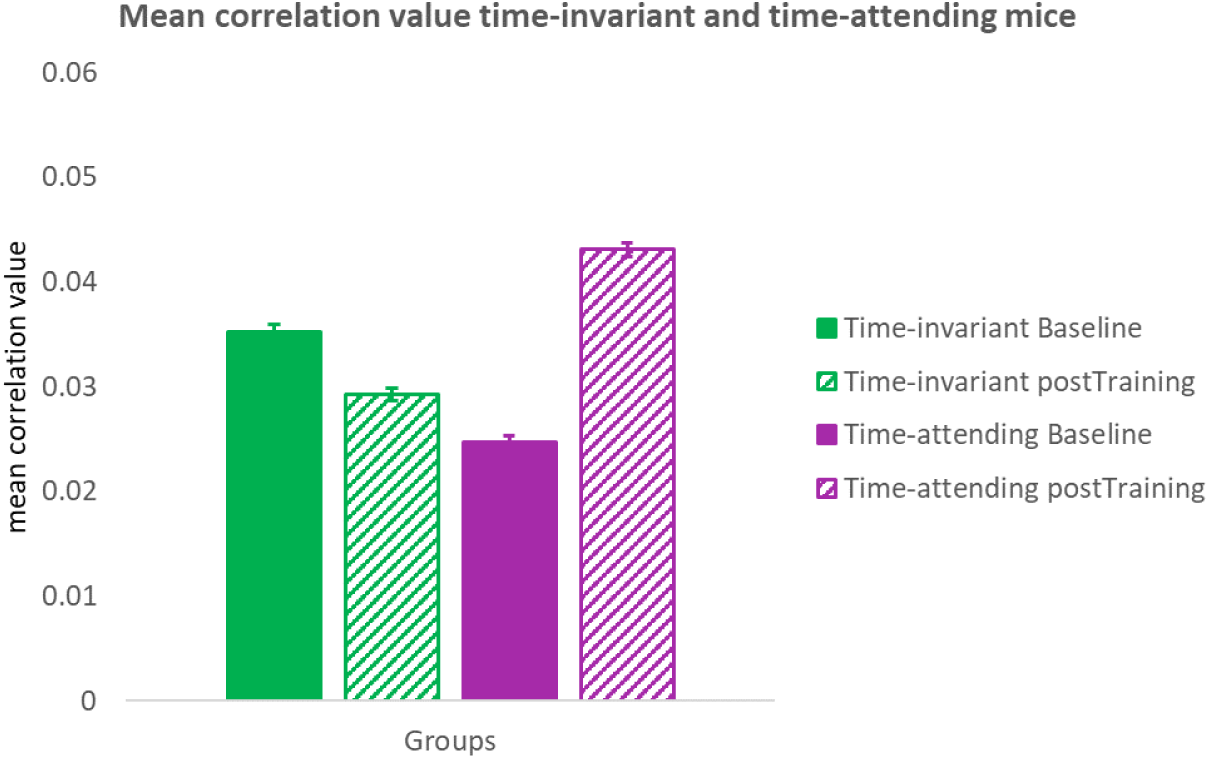
Mean correlation values of time-invariant and time-attending mice based on correlation analysis of the resting-state fMRI. The mean correlation coefficient and the corresponding s.e.m. were calculated for both groups of animals (time-invariant [green], time-attending [purple]) at baseline (filled) and post training (shaded) based on their correlation values extracted from the individual Pearson correlation matrix.

**Supplementary Figure 6.**
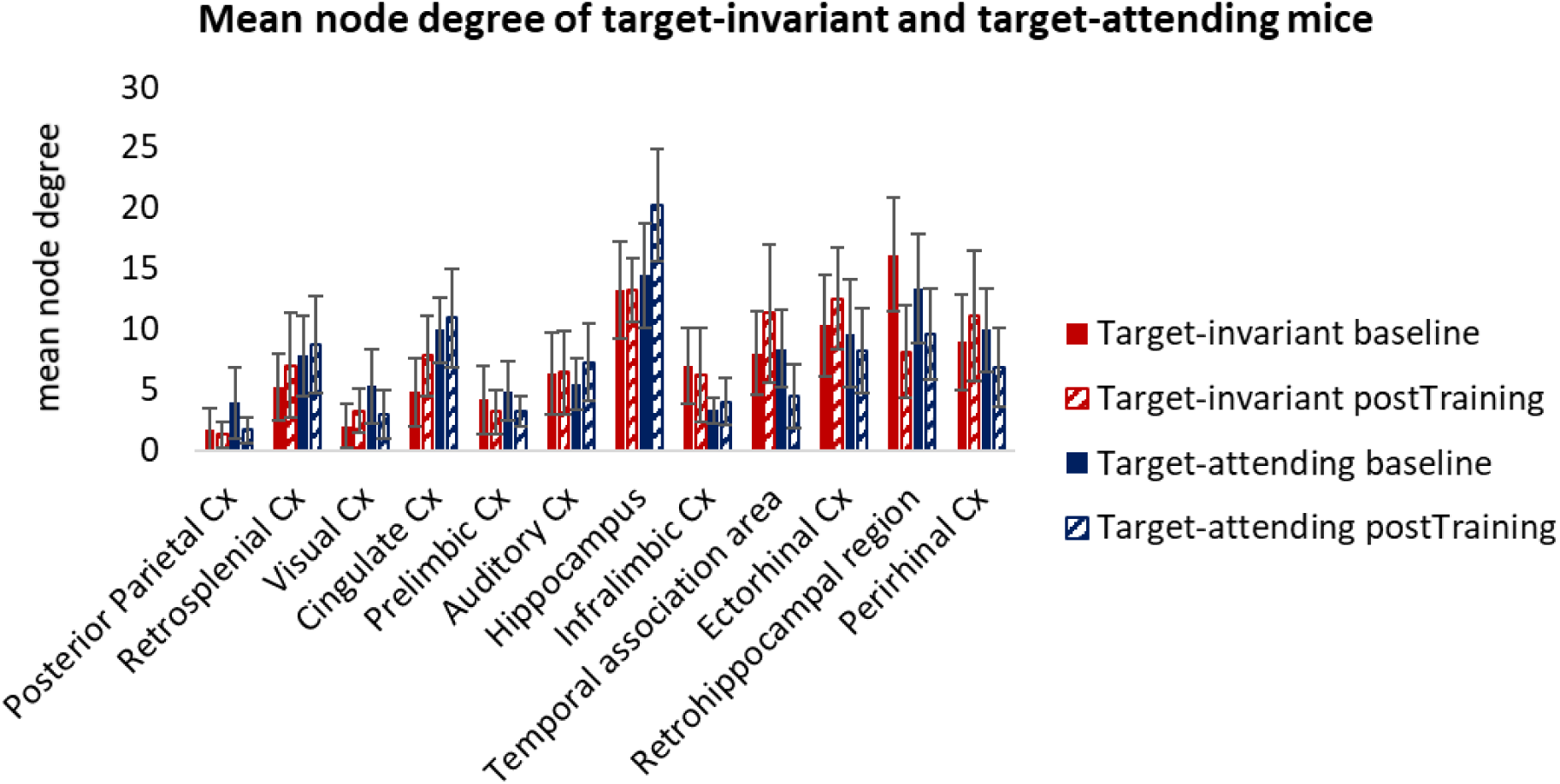
Mean node degree of target-invariant and target-attending mice based on graph-theory analyses of the resting-state fMRI for selected regions of interest. The mean node degree and the corresponding s.e.m. were calculated for both groups of animals (target-invariant [red], target-attending [blue]) at baseline (filled) and post training (shaded) based on their extracted individual node degree from a graph-theoretical approach. Values from the right and left hemisphere were averaged to simplify visualization and interpretation.

**Supplementary Figure 7.**
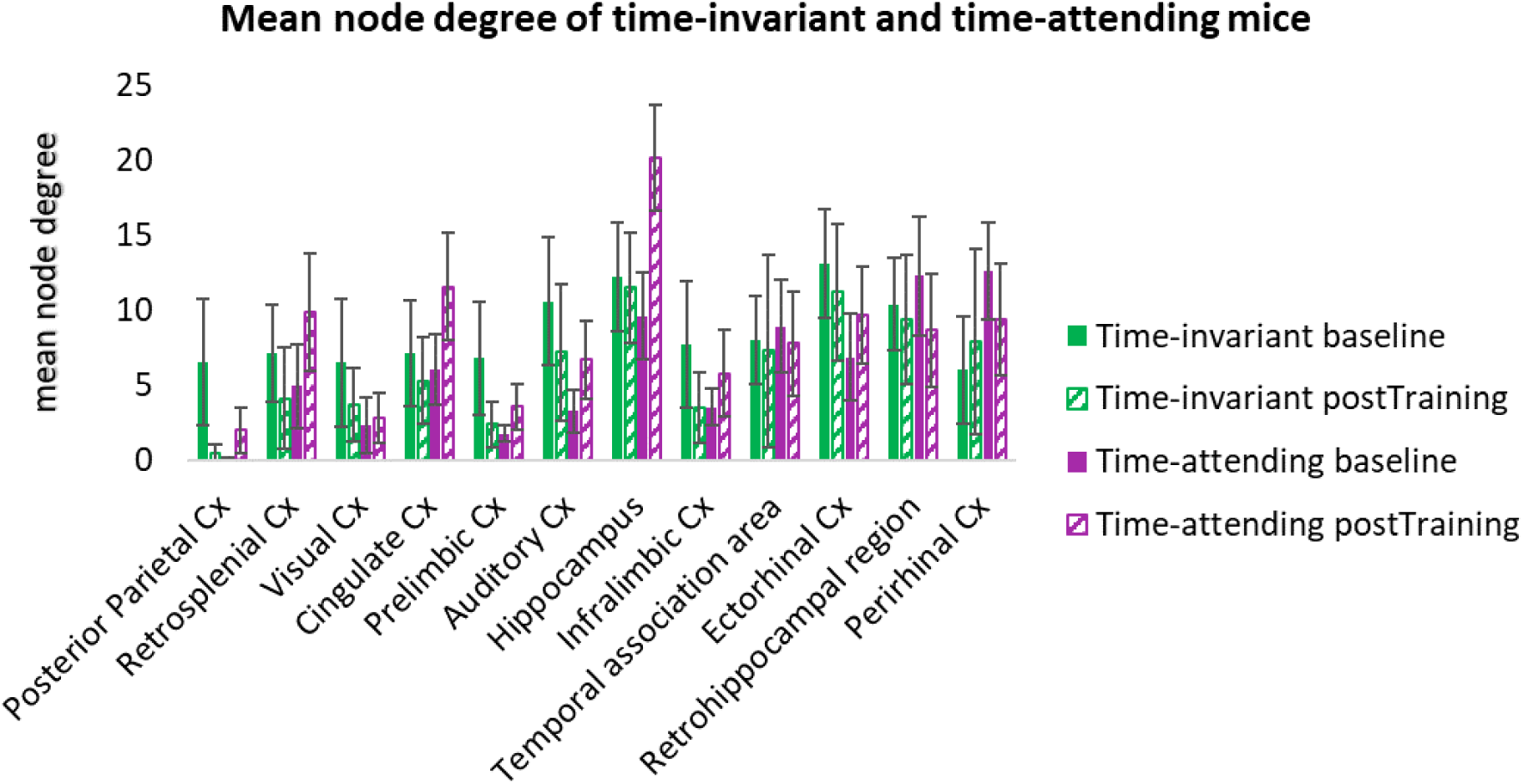
Mean node degree of time-invariant and time-attending mice based on graph-theory analyses of the resting-state fMRI for selected regions of interest. The mean node degree and the corresponding s.e.m. were calculated for both groups of animals (time-invariant [green], time-attending [purple]) at baseline (filled) and post training (shaded) based on their extracted individual node degree from a graph-theoretical approach. Values from the right and left hemisphere were averaged to simplify visualization and interpretation.

## References

Adams, J. N., Maass, A., & Berron, D. (2021). Reduced repetition suppression in aging is driven by tau–related hyperactivity in medial temporal lobe. Journal of. https://www.jneurosci.org/content/41/17/3917.abstract

Alwashmi, K., Meyer, G., Rowe, F., & Ward, R. (2024). Enhancing learning outcomes through multisensory integration: A fMRI study of audio-visual training in virtual reality. NeuroImage, 285, 120483. 10.1016/j.neuroimage.2023.120483

Ashwood, Z. C., Roy, N. A., Stone, I. R., International Brain Laboratory, Urai, A. E., Churchland, A. K., Pouget, A., & Pillow, J. W. (2022). Mice alternate between discrete strategies during perceptual decision-making. Nature Neuroscience, 25(2), 201–212. 10.1038/s41593-021-01007-z

Balasubramaniam, R., Haegens, S., Jazayeri, M., Merchant, H., Sternad, D., & Song, J.-H. (2021). Neural Encoding and Representation of Time for Sensorimotor Control and Learning. The Journal of Neuroscience: The Official Journal of the Society for Neuroscience, 41(5), 866–872. 10.1523/JNEUROSCI.1652-20.2020

Ball, F., Andreca, J., & Noesselt, T. (2022). Context dependency of time-based event-related expectations for different modalities. Psychological Research, 86(4), 1239–1251. 10.1007/s00426-021-01564-9

Ball, F., Fuehrmann, F., Stratil, F., & Noesselt, T. (2018). Phasic and sustained interactions of multisensory interplay and temporal expectation. Scientific Reports, 8(1), 10208. 10.1038/s41598-018-28495-7

Ball, F., Michels, L. E., Thiele, C., & Noesselt, T. (2018). The role of multisensory interplay in enabling temporal expectations. Cognition, 170, 130–146. 10.1016/j.cognition.2017.09.015

Bentley, W. J., Li, J. M., Snyder, A. Z., Raichle, M. E., & Snyder, L. H. (2016). Oxygen Level and LFP in Task-Positive and Task-Negative Areas: Bridging BOLD fMRI and Electrophysiology. Cerebral Cortex, 26(1), 346–357. 10.1093/cercor/bhu260

Billette, O. V., Ziegler, G., Aruci, M., Schütze, H., Kizilirmak, J. M., Richter, A., Altenstein, S., Bartels, C., Brosseron, F., Cardenas-Blanco, A., Dahmen, P., Dechent, P., Dobisch, L., Fliessbach, K., Freiesleben, S. D., Glanz, W., Göerß, D., Haynes, J. D., Heneka, M. T., … DELCODE Study Group. (2022). Novelty-Related fMRI Responses of Precuneus and Medial Temporal Regions in Individuals at Risk for Alzheimer Disease. Neurology, 99(8), e775–e788. 10.1212/WNL.0000000000200667

Biswal, B., Yetkin, F. Z., Haughton, V. M., & Hyde, J. S. (1995). Functional connectivity in the motor cortex of resting human brain using echo-planar MRI. Magnetic Resonance in Medicine: Official Journal of the Society of Magnetic Resonance in Medicine / Society of Magnetic Resonance in Medicine, 34(4), 537–541. 10.1002/mrm.1910340409

Bressler, D., Spotswood, N., & Whitney, D. (2007). Negative BOLD fMRI response in the visual cortex carries precise stimulus-specific information. PloS One, 2(5), e410. 10.1371/journal.pone.0000410

Buzsáki, G., & Llinás, R. (2017). Space and time in the brain. Science, 358(6362), 482–485. 10.1126/science.aan8869

Caselli, L., Iaboli, L., & Nichelli, P. (2009). Time estimation in mild Alzheimer’s disease patients. Behavioral and Brain Functions: BBF, 5(1), 32. 10.1186/1744-9081-5-32

Chapuis, G. A., & Chadderton, P. T. (2018). Using temporal expectation to assess auditory streaming in mice. Frontiers in Behavioral Neuroscience, 12, 205. 10.3389/fnbeh.2018.00205

Choi, U.-S., Sung, Y.-W., & Ogawa, S. (2017). Steady-state and dynamic network modes for perceptual expectation. Scientific Reports, 7, 40626. 10.1038/srep40626

De Corte, B. J., Farley, S. J., Heslin, K. A., Parker, K. L., & Freeman, J. H. (2022). The dorsal hippocampus’ role in context-based timing in rodents. Neurobiology of Learning and Memory, 194, 107673. 10.1016/j.nlm.2022.107673

Diersch, N., Valdes-Herrera, J. P., Tempelmann, C., & Wolbers, T. (2021). Increased Hippocampal Excitability and Altered Learning Dynamics Mediate Cognitive Mapping Deficits in Human Aging. The Journal of Neuroscience: The Official Journal of the Society for Neuroscience, 41(14), 3204– 3221. 10.1523/JNEUROSCI.0528-20.2021

Driver, J., & Noesselt, T. (2008). Multisensory interplay reveals crossmodal influences on “sensory-specific” brain regions, neural responses, and judgments. Neuron, 57, 11–23. 10.1016/j.neuron.2007.12.013

Dylda, E., & Pakan, J. M. P. (2021). Visual plasticity: Illuminating the role of the hippocampus in cortical sensory encoding [Review of *Visual plasticity: Illuminating the role of the hippocampus in cortical sensory encoding*]. Current Biology: CB, 31(18), R1087–R1089. 10.1016/j.cub.2021.07.015

Dylda, E., Pakan, J. M. P., & Rochefort, N. L. (2018). Chapter 13 - Chronic Two-Photon Calcium Imaging in the Visual Cortex of Awake Behaving Mice. In D. Manahan-Vaughan (Ed.), Handbook of Behavioral Neuroscience (Vol. 28, pp. 235–251). Elsevier. 10.1016/B978-0-12-812028-6.00013-6

El Haj, M., & Kapogiannis, D. (2016). Time distortions in Alzheimer’s disease: a systematic review and theoretical integration. NPJ Aging and Mechanisms of Disease, 2, 16016. 10.1038/npjamd.2016.16

Ernst, M. O., & Bülthoff, H. H. (2004). Merging the senses into a robust percept. Trends in Cognitive Sciences, 8(4), 162–169. 10.1016/j.tics.2004.02.002

Fiehler, K., Brenner, E., & Spering, M. (2019). Prediction in goal-directed action. Journal of Vision, 19(9), 10. 10.1167/19.9.10

Finnerty, G. T., Shadlen, M. N., Jazayeri, M., Nobre, A. C., & Buonomano, D. V. (2015). Time in Cortical Circuits. The Journal of Neuroscience: The Official Journal of the Society for Neuroscience, 35(41), 13912–13916. 10.1523/JNEUROSCI.2654-15.2015

Finnie, P. S. B., Komorowski, R. W., & Bear, M. F. (2021). The spatiotemporal organization of experience dictates hippocampal involvement in primary visual cortical plasticity. Current Biology: CB, 31(18), 3996–4008.e6. 10.1016/j.cub.2021.06.079

Garner, A. R., & Keller, G. B. (2022). A cortical circuit for audio-visual predictions. Nature Neuroscience, 25(1), 98–105. 10.1038/s41593-021-00974-7

Gazzaley, A., & Nobre, A. C. (2012). Top-down modulation: bridging selective attention and working memory. Trends in Cognitive Sciences, 16(2), 129–135. 10.1016/j.tics.2011.11.014

Gil, R., Valente, M., & Shemesh, N. (2024). Rat superior colliculus encodes the transition between static and dynamic vision modes. Nature Communications, 15(1), 849. 10.1038/s41467-024-44934-8

Hawellek, D. J., Wong, Y. T., & Pesaran, B. (2016). Temporal coding of reward-guided choice in the posterior parietal cortex. Proceedings of the National Academy of Sciences of the United States of America, 113(47), 13492–13497. 10.1073/pnas.1606479113

He, H., Ettehadi, N., Shmuel, A., & Razlighi, Q. R. (2022). Evidence suggesting common mechanisms underlie contralateral and ipsilateral negative BOLD responses in the human visual cortex. NeuroImage, 262, 119440. 10.1016/j.neuroimage.2022.119440

Henschke, J. U., Dylda, E., Katsanevaki, D., Dupuy, N., Currie, S. P., Amvrosiadis, T., Pakan, J. M. P., & Rochefort, N. L. (2020). Reward Association Enhances Stimulus-Specific Representations in Primary Visual Cortex. Current Biology: CB, 30(10), 1866–1880.e5. 10.1016/j.cub.2020.03.018

Heys, J. G., & Dombeck, D. A. (2018). Evidence for a subcircuit in medial entorhinal cortex representing elapsed time during immobility. Nature Neuroscience, 21(11), 1574–1582. 10.1038/s41593-018-0252-8

Holtmaat, A., Bonhoeffer, T., Chow, D. K., Chuckowree, J., De Paola, V., Hofer, S. B., Hübener, M., Keck, T., Knott, G., Lee, W.-C. A., Mostany, R., Mrsic-Flogel, T. D., Nedivi, E., Portera-Cailliau, C., Svoboda, K., Trachtenberg, J. T., & Wilbrecht, L. (2009). Long-term, high-resolution imaging in the mouse neocortex through a chronic cranial window. Nature Protocols, 4(8), 1128–1144. 10.1038/nprot.2009.89

Huang, W., Wang, Y., Qin, J., He, C., Li, Y., Wang, Y., Li, M., Lyu, J., Zhou, Z., Jia, H., Pakan, J., Xie, P., & Zhang, J. (2023). A corticostriatal projection for sound-evoked and anticipatory motor behavior following temporal expectation. Neuroreport, 34(1), 1–8. 10.1097/WNR.0000000000001851

Hu, D., & Huang, L. (2015). Negative hemodynamic response in the cortex: evidence opposing neuronal deactivation revealed via optical imaging and electrophysiological recording. Journal of Neurophysiology, 114(4), 2152–2161. 10.1152/jn.00246.2015

Jaramillo, S., & Zador, A. M. (2011). The auditory cortex mediates the perceptual effects of acoustic temporal expectation. Nature Neuroscience, 14(2), 246–251. 10.1038/nn.2688

Jenkinson, M., Bannister, P., Brady, M., & Smith, S. (2002). Improved optimization for the robust and accurate linear registration and motion correction of brain images. NeuroImage, 17(2), 825–841. 10.1016/s1053-8119(02)91132-8

Jenkinson, M., Beckmann, C. F., Behrens, T. E. J., Woolrich, M. W., & Smith, S. M. (2012). FSL. NeuroImage, 62(2), 782–790. 10.1016/j.neuroimage.2011.09.015

Jenkinson, M., & Smith, S. (2001). A global optimisation method for robust affine registration of brain images. Medical Image Analysis, 5(2), 143–156. 10.1016/s1361-8415(01)00036-6

Jin, W., Nobre, A. C., & van Ede, F. (2020). Temporal Expectations Prepare Visual Working Memory for Behavior. Journal of Cognitive Neuroscience, 32(12), 2320–2332. 10.1162/jocn_a_01626

Jones, A., Ward, E. V., Csiszer, E. L., & Szymczak, J. (2022). Temporal Expectation Improves Recognition Memory for Spatially Attended Objects. Journal of Cognitive Neuroscience, 34(9), 1616–1629. 10.1162/jocn_a_01872

Kaifosh, P., Zaremba, J. D., Danielson, N. B., & Losonczy, A. (2014). SIMA: Python software for analysis of dynamic fluorescence imaging data. Frontiers in Neuroinformatics, 8, 80. 10.3389/fninf.2014.00080

Keemink, S. W., Lowe, S. C., Pakan, J. M. P., Dylda, E., van Rossum, M. C. W., & Rochefort, N. L. (2018). FISSA: A neuropil decontamination toolbox for calcium imaging signals. Scientific Reports, 8(1), 3493. 10.1038/s41598-018-21640-2

Keller, G. B., & Mrsic-Flogel, T. D. (2018). Predictive Processing: A Canonical Cortical Computation. Neuron, 100(2), 424–435. 10.1016/j.neuron.2018.10.003

Knudstrup, S. G., Martinez, C., & Gavornik, J. P. (2024). Learned response dynamics reflect stimulus timing and encode temporal expectation violations in superficial layers of mouse V1. In eLife. 10.7554/elife.94727.1

Kraus, B. J., Robinson, R. J., 2nd, White, J. A., Eichenbaum, H., & Hasselmo, M. E. (2013). Hippocampal “time cells”: time versus path integration. Neuron, 78(6), 1090–1101. 10.1016/j.neuron.2013.04.015

Lambers, H., Segeroth, M., Albers, F., Wachsmuth, L., van Alst, T. M., & Faber, C. (2020). A cortical rat hemodynamic response function for improved detection of BOLD activation under common experimental conditions. NeuroImage, 208, 116446. 10.1016/j.neuroimage.2019.116446

Le, N. M., Yildirim, M., Wang, Y., Sugihara, H., Jazayeri, M., & Sur, M. (2023). Mixtures of strategies underlie rodent behavior during reversal learning. PLoS Computational Biology, 19(9), e1011430. 10.1371/journal.pcbi.1011430

Li, J., Liao, X., Zhang, J., Wang, M., Yang, N., Zhang, J., Lv, G., Li, H., Lu, J., Ding, R., Li, X., Guang, Y., Yang, Z., Qin, H., Jin, W., Zhang, K., He, C., Jia, H., Zeng, S., … Chen, X. (2017). Primary Auditory Cortex is Required for Anticipatory Motor Response. Cerebral Cortex, 27(6), 3254–3271. 10.1093/cercor/bhx079

Li, R., Huang, J., Li, L., Zhao, Z., Liang, S., Liang, S., Wang, M., Liao, X., Lyu, J., Zhou, Z., Wang, S., Jin, W., Chen, H., Holder, D., Liu, H., Zhang, J., Li, M., Tang, Y., Remy, S., … Jia, H. (2023). Holistic bursting cells store long-term memory in auditory cortex. Nature Communications, 14(1), 8090. 10.1038/s41467-023-43620-5

Livesey, A. C., Wall, M. B., & Smith, A. T. (2007). Time perception: manipulation of task difficulty dissociates clock functions from other cognitive demands. Neuropsychologia, 45(2), 321–331. 10.1016/j.neuropsychologia.2006.06.033

Logothetis, N. K., Pauls, J., Augath, M., Trinath, T., & Oeltermann, A. (2001). Neurophysiological investigation of the basis of the fMRI signal. Nature, 412(6843), 150–157. 10.1038/35084005

Lyamzin, D., & Benucci, A. (2019). The mouse posterior parietal cortex: Anatomy and functions. Neuroscience Research, 140, 14–22. 10.1016/j.neures.2018.10.008

MacDonald, C. J., Lepage, K. Q., Eden, U. T., & Eichenbaum, H. (2011). Hippocampal “time cells” bridge the gap in memory for discontiguous events. Neuron, 71(4), 737–749. 10.1016/j.neuron.2011.07.012

Malhotra, P., Coulthard, E. J., & Husain, M. (2009). Role of right posterior parietal cortex in maintaining attention to spatial locations over time. Brain: A Journal of Neurology, 132(Pt 3), 645– 660. 10.1093/brain/awn350

Marek, S., & Dosenbach, N. U. F. (2018). The frontoparietal network: function, electrophysiology, and importance of individual precision mapping. Dialogues in Clinical Neuroscience, 20(2), 133–140. 10.31887/DCNS.2018.20.2/smarek

Mathis, A., Mamidanna, P., Cury, K. M., Abe, T., Murthy, V. N., Mathis, M. W., & Bethge, M. (2018). DeepLabCut: markerless pose estimation of user-defined body parts with deep learning. Nature Neuroscience, 21(9), 1281–1289. 10.1038/s41593-018-0209-y

Ma, Y., Shaik, M. A., Kozberg, M. G., Kim, S. H., Portes, J. P., Timerman, D., & Hillman, E. M. C. (2016). Resting-state hemodynamics are spatiotemporally coupled to synchronized and symmetric neural activity in excitatory neurons. Proceedings of the National Academy of Sciences of the United States of America, 113(52), E8463–E8471. 10.1073/pnas.1525369113

Modat, M., Cash, D. M., Daga, P., Winston, G. P., Duncan, J. S., & Ourselin, S. (2014). Global image registration using a symmetric block-matching approach. Journal of Medical Imaging (Bellingham, Wash.), 1(2), 024003. 10.1117/1.JMI.1.2.024003

Modat, M., Ridgway, G. R., Taylor, Z. A., Lehmann, M., Barnes, J., Hawkes, D. J., Fox, N. C., & Ourselin, S. (2010). Fast free-form deformation using graphics processing units. Computer Methods and Programs in Biomedicine, 98(3), 278–284. 10.1016/j.cmpb.2009.09.002

Montijn, J. S., Vinck, M., & Pennartz, C. M. A. (2014). Population coding in mouse visual cortex: response reliability and dissociability of stimulus tuning and noise correlation. Frontiers in Computational Neuroscience, 8, 58. 10.3389/fncom.2014.00058

Narayanan, N. S., & Laubach, M. (2009). Delay activity in rodent frontal cortex during a simple reaction time task. Journal of Neurophysiology, 101(6), 2859–2871. 10.1152/jn.90615.2008

Nobre, A. C., & van Ede, F. (2018). Anticipated moments: temporal structure in attention. Nature Reviews. Neuroscience, 19(1), 34–48. 10.1038/nrn.2017.141

Ourselin, S., Roche, A., Subsol, G., Pennec, X., & Ayache, N. (2001). Reconstructing a 3D structure from serial histological sections. Image and Vision Computing, 19(1), 25–31. 10.1016/S0262-8856(00)00052-4

Pakan, J. M. P., Currie, S. P., Fischer, L., & Rochefort, N. L. (2018). The Impact of Visual Cues, Reward, and Motor Feedback on the Representation of Behaviorally Relevant Spatial Locations in Primary Visual Cortex. Cell Reports, 24(10), 2521–2528. 10.1016/j.celrep.2018.08.010

Pakan, J. M. P., Francioni, V., & Rochefort, N. L. (2018). Action and learning shape the activity of neuronal circuits in the visual cortex. Current Opinion in Neurobiology, 52, 88–97. 10.1016/j.conb.2018.04.020

Pakan, J. M. P., Lowe, S. C., Dylda, E., Keemink, S. W., Currie, S. P., Coutts, C. A., & Rochefort, N. L. (2016). Behavioral-state modulation of inhibition is context-dependent and cell type specific in mouse visual cortex. eLife, 5. 10.7554/eLife.14985

Pallast, N., Diedenhofen, M., Blaschke, S., Wieters, F., Wiedermann, D., Hoehn, M., Fink, G. R., & Aswendt, M. (2019). Processing Pipeline for Atlas-Based Imaging Data Analysis of Structural and Functional Mouse Brain MRI (AIDAmri). Frontiers in Neuroinformatics, 13, 42. 10.3389/fninf.2019.00042

Pastalkova, E., Itskov, V., Amarasingham, A., & Buzsáki, G. (2008). Internally generated cell assembly sequences in the rat hippocampus. Science, 321(5894), 1322–1327. 10.1126/science.1159775

Pinto, L., Rajan, K., DePasquale, B., Thiberge, S. Y., Tank, D. W., & Brody, C. D. (2019). Task-Dependent Changes in the Large-Scale Dynamics and Necessity of Cortical Regions. Neuron, 104(4), 810–824.e9. 10.1016/j.neuron.2019.08.025

Ripp, I., Zur Nieden, A.-N., Blankenagel, S., Franzmeier, N., Lundström, J. N., & Freiherr, J. (2018). Multisensory integration processing during olfactory-visual stimulation-An fMRI graph theoretical network analysis. Human Brain Mapping, 39(9), 3713–3727. 10.1002/hbm.24206

Rohenkohl, G., Cravo, A. M., Wyart, V., & Nobre, A. C. (2012). Temporal Expectation Improves the Quality of Sensory Information. The Journal of Neuroscience: The Official Journal of the Society for Neuroscience, 32(24), 8424–8428. 10.1523/JNEUROSCI.0804-12.2012

Roy, N. A., Bak, J. H., Akrami, A., Brody, C. D., & Pillow, J. W. (2018). Efficient inference for time-varying behavior during learning. Advances in Neural Information Processing Systems, 31, 5695– 5705. https://www.ncbi.nlm.nih.gov/pubmed/31244514

Rubinov, M., & Sporns, O. (2010). Complex network measures of brain connectivity: uses and interpretations. NeuroImage, 52(3), 1059–1069. 10.1016/j.neuroimage.2009.10.003

Rueckert, D., Sonoda, L. I., Hayes, C., Hill, D. L., Leach, M. O., & Hawkes, D. J. (1999). Nonrigid registration using free-form deformations: application to breast MR images. IEEE Transactions on Medical Imaging, 18(8), 712–721. 10.1109/42.796284

Scharwächter, L., Schmitt, F. J., Pallast, N., Fink, G. R., & Aswendt, M. (2022). Network analysis of neuroimaging in mice. NeuroImage, 253, 119110. 10.1016/j.neuroimage.2022.119110

Schridde, U., Khubchandani, M., Motelow, J. E., Sanganahalli, B. G., Hyder, F., & Blumenfeld, H. (2008). Negative BOLD with large increases in neuronal activity. Cerebral Cortex, 18(8), 1814– 1827. 10.1093/cercor/bhm208

Schubotz, R. I. (2007). Prediction of external events with our motor system: towards a new framework. Trends in Cognitive Sciences, 11(5), 211–218. 10.1016/j.tics.2007.02.006

Sharma, J., Sugihara, H., Katz, Y., Schummers, J., Tenenbaum, J., & Sur, M. (2015). Spatial Attention and Temporal Expectation Under Timed Uncertainty Predictably Modulate Neuronal Responses in Monkey V1. Cerebral Cortex, 25(9), 2894–2906. 10.1093/cercor/bhu086

Shmuel, A., Augath, M., Oeltermann, A., & Logothetis, N. K. (2006). Negative functional MRI response correlates with decreases in neuronal activity in monkey visual area V1. Nature Neuroscience, 9(4), 569–577. 10.1038/nn1675

Shuler, M. G. H. (2016). Timing in the visual cortex and its investigation. Current Opinion in Behavioral Sciences, 8, 73–77. 10.1016/j.cobeha.2016.02.006

Smith, A. J., Blumenfeld, H., Behar, K. L., Rothman, D. L., Shulman, R. G., & Hyder, F. (2002). Cerebral energetics and spiking frequency: the neurophysiological basis of fMRI. Proceedings of the National Academy of Sciences of the United States of America, 99(16), 10765–10770. 10.1073/pnas.132272199

Smith, S. M. (2002). Fast robust automated brain extraction. Human Brain Mapping, 17(3), 143–155. 10.1002/hbm.10062

Smith, S. M., Jenkinson, M., Woolrich, M. W., Beckmann, C. F., Behrens, T. E. J., Johansen-Berg, H., Bannister, P. R., De Luca, M., Drobnjak, I., Flitney, D. E., Niazy, R. K., Saunders, J., Vickers, J., Zhang, Y., De Stefano, N., Brady, J. M., & Matthews, P. M. (2004). Advances in functional and structural MR image analysis and implementation as FSL. NeuroImage, 23 *Suppl 1*, S208–S219. 10.1016/j.neuroimage.2004.07.051

Stefanics, G., Hangya, B., Hernádi, I., Winkler, I., Lakatos, P., & Ulbert, I. (2010). Phase entrainment of human delta oscillations can mediate the effects of expectation on reaction speed. The Journal of Neuroscience: The Official Journal of the Society for Neuroscience, 30(41), 13578–13585. 10.1523/JNEUROSCI.0703-10.2010

Stefanovic, B., Warnking, J. M., & Pike, G. B. (2004). Hemodynamic and metabolic responses to neuronal inhibition. NeuroImage, 22(2), 771–778. 10.1016/j.neuroimage.2004.01.036

Subramanian, D. L., & Smith, D. M. (2024). Time Cells in the Retrosplenial Cortex. In bioRxiv (p. 2024.03.01.583039). 10.1101/2024.03.01.583039

Sullender, C. T., Richards, L. M., He, F., Luan, L., & Dunn, A. K. (2022). Dynamics of isoflurane-induced vasodilation and blood flow of cerebral vasculature revealed by multi-exposure speckle imaging. Journal of Neuroscience Methods, 366, 109434. 10.1016/j.jneumeth.2021.109434

Sun, W., Choi, I., Stoyanov, S., Senkov, O., Ponimaskin, E., Winter, Y., Pakan, J. M. P., & Dityatev, A. (2021). Context value updating and multidimensional neuronal encoding in the retrosplenial cortex. Nature Communications, 12(1), 6045. 10.1038/s41467-021-26301-z

Tang, M., Kheradpezhouh, E., Lee, C. C. Y., Dickinson, J. E., Mattingley, J. B., & Arabzadeh, E. (2023). Expectation violations enhance neuronal encoding of sensory information in mouse primary visual cortex. Nature Communications, 14(1), 1196. 10.1038/s41467-023-36608-8

Tang, X., Wu, J., & Shen, Y. (2016). The interactions of multisensory integration with endogenous and exogenous attention. Neuroscience and Biobehavioral Reviews, 61, 208–224. 10.1016/j.neubiorev.2015.11.002

Tavares, T. F., Bueno, J. L. O., & Doyère, V. (2022). Temporal prediction error triggers amygdala-dependent memory updating in appetitive operant conditioning in rats. Frontiers in Behavioral Neuroscience, 16, 1060587. 10.3389/fnbeh.2022.1060587

Taxidis, J., Pnevmatikakis, E. A., Dorian, C. C., Mylavarapu, A. L., Arora, J. S., Samadian, K. D., Hoffberg, E. A., & Golshani, P. (2020). Differential Emergence and Stability of Sensory and Temporal Representations in Context-Specific Hippocampal Sequences. Neuron, 108(5), 984– 998.e9. 10.1016/j.neuron.2020.08.028

Terada, S., Sakurai, Y., Nakahara, H., & Fujisawa, S. (2017). Temporal and Rate Coding for Discrete Event Sequences in the Hippocampus. Neuron, 94(6), 1248–1262.e4. 10.1016/j.neuron.2017.05.024

Todd, T. P., Fournier, D. I., & Bucci, D. J. (2019). Retrosplenial cortex and its role in cue-specific learning and memory. Neuroscience and Biobehavioral Reviews, 107, 713–728. 10.1016/j.neubiorev.2019.04.016

Urai, A. E., Aguillon-Rodriguez, V., Laranjeira, I. C., Cazettes, F., International Brain Laboratory, Mainen, Z. F., & Churchland, A. K. (2021). Citric Acid Water as an Alternative to Water Restriction for High-Yield Mouse Behavior. eNeuro, 8(1). 10.1523/ENEURO.0230-20.2020

Van Aken, H., Fitch, W., Graham, D. I., Brüssel, T., & Themann, H. (1986). Cardiovascular and cerebrovascular effects of isoflurane-induced hypotension in the baboon. Anesthesia and Analgesia, 65(6), 565–574. https://www.ncbi.nlm.nih.gov/pubmed/3706797

van Duuren, E., Lankelma, J., & Pennartz, C. M. A. (2008). Population coding of reward magnitude in the orbitofrontal cortex of the rat. The Journal of Neuroscience: The Official Journal of the Society for Neuroscience, 28(34), 8590–8603. 10.1523/JNEUROSCI.5549-07.2008

Wang, L., & Krauzlis, R. J. (2018). Visual Selective Attention in Mice. Current Biology: CB, 28(5), 676–685.e4. 10.1016/j.cub.2018.01.038

Wang, M., Liao, X., Li, R., Liang, S., Ding, R., Li, J., Zhang, J., He, W., Liu, K., Pan, J., Zhao, Z., Li, T., Zhang, K., Li, X., Lyu, J., Zhou, Z., Varga, Z., Mi, Y., Zhou, Y., … Chen, X. (2020). Single-neuron representation of learned complex sounds in the auditory cortex. Nature Communications, 11(1), 4361. 10.1038/s41467-020-18142-z

Wang, Q., Ding, S.-L., Li, Y., Royall, J., Feng, D., Lesnar, P., Graddis, N., Naeemi, M., Facer, B., Ho, A., Dolbeare, T., Blanchard, B., Dee, N., Wakeman, W., Hirokawa, K. E., Szafer, A., Sunkin, S. M., Oh, S. W., Bernard, A., … Ng, L. (2020). The Allen Mouse Brain Common Coordinate Framework: A 3D Reference Atlas. Cell, 181(4), 936–953.e20. 10.1016/j.cell.2020.04.007

Whitlock, J. R. (2017). Posterior parietal cortex. Current Biology: CB, 27(14), R691–R695. 10.1016/j.cub.2017.06.007

Woolrich, M. W., Jbabdi, S., Patenaude, B., Chappell, M., Makni, S., Behrens, T., Beckmann, C., Jenkinson, M., & Smith, S. M. (2009). Bayesian analysis of neuroimaging data in FSL. NeuroImage, *45*(1 Suppl), S173–S186. 10.1016/j.neuroimage.2008.10.055

Zhang, Y., Brady, M., & Smith, S. (2001). Segmentation of brain MR images through a hidden Markov random field model and the expectation-maximization algorithm. IEEE Transactions on Medical Imaging, 20(1), 45–57. 10.1109/42.906424

Zierul, B., Röder, B., Tempelmann, C., Bruns, P., & Noesselt, T. (2017). The role of auditory cortex in the spatial ventriloquism aftereffect. NeuroImage, 162, 257–268. 10.1016/j.neuroimage.2017.09.002

